# A genetic framework for RNAi inheritance in *Caenorhabditis elegans*

**DOI:** 10.1101/2024.10.02.616260

**Authors:** Jan Schreier, Fridolin Kielisch, René F. Ketting

## Abstract

Gene regulation by RNA interference (RNAi) is a conserved process driven by double-stranded RNA (dsRNA). It responds to exogenous cues and drives endogenous gene regulation. In *Caenorhabditis elegans*, RNAi can be inherited from parents to offspring. While a number of factors have been implicated in this inheritance process, we do not understand how and when they function. Using a new inheritance assay, we establish a hierarchy amongst previously identified inheritance factors. The nuclear argonaute protein HRDE-1 was required for RNAi establishment in parents and offspring, but not for the inheritance process. In contrast, the cytoplasmic argonaute protein WAGO-3 was the only factor essential for inheritance, via sperm and oocyte, while not affecting establishment in either parent or offspring. We propose a cycle between nuclear and cytoplasmic argonaute proteins, where nuclear activity drives most of the silencing and cytoplasmic activity ensures inheritance. Finally, we implicate the RNA helicase ZNFX-1 as a factor that controls the entry of exogenous versus endogenous small RNAs into this cycle, ensuring a proper balance between gene silencing and activity.

## INTRODUCTION

RNA interference (RNAi) is a process in which genes are silenced through double-stranded RNA (dsRNA)(Fire *et al*, 1998). After processing the dsRNA into smaller fragments by the enzyme Dicer, these so-called siRNAs are bound by a protein of the Argonaute family. Guided by siRNAs, these proteins can induce the destabilisation of mRNAs by direct cleavage or through the recruitment of additional factors. Additionally, nuclear Argonaute proteins can induce transcriptional gene silencing. These processes, reviewed by (Ketting, 2011; Meister, 2013; Ozata *et al*, 2019; Luteijn & Ketting, 2013), or similar processes are deeply conserved and they play a role various aspects of gene activity control.

In *Caenorhabditis elegans*, the RNAi process is driven through two-step process (reviewed in (Ketting & Cochella, 2021)). First, the siRNAs made from the dsRNA are bound by an Argonaute protein named RDE-1. Instead of directly silencing mRNAs with complementary sequences, RDE-1 triggers the activity of an RNA-dependent RNA polymerase (RRF-1) on the targeted mRNA. RRF-1 generates secondary siRNAs, also known as 22G RNAs, which are bound by worm-specific Argonaute proteins, or WAGOs. These then drive the silencing in an as yet poorly understood manner. This amplification step also allows for another feature: the inheritance of RNAi-driven silencing across generations (Grishok *et al*, 2000; Fire *et al*, 1998; Alcazar *et al*, 2008; Vastenhouw *et al*, 2006). In many cases silencing up to six generations after the dsRNA-exposed generation can still be observed, and in some cases, virtually indefinitely stable silencing can be induced, albeit that this has been observed only in strains lacking the Argonaute protein PRG-1 (Luteijn *et al*, 2012; Shirayama *et al*, 2012; Ashe *et al*, 2012; Shukla *et al*, 2021). Normally, RNAi inheritance is restricted in duration, and specific mechanisms may be in place to ensure that inheritance is finite (Houri-Ze’evi *et al*, 2016).

The combination of environmental effects on gene expression through dsRNA and the inheritance of this silencing over generations has led to studies probing whether environmental cues, in the form of gene regulatory information, may be transmitted to offspring. Potentially, this may better prepare animals to face specific environmental conditions that were already met by their parents. Indeed, links between nutrients (Rechavi *et al*, 2014), pathogens (Rechavi *et al*, 2011; Sengupta *et al*, 2024; Kaletsky *et al*, 2020) and animal behaviour (Posner *et al*, 2019; Toker *et al*, 2022) have been coupled to RNAi inheritance, although in some cases there is dispute on reproducibility (Gainey *et al*, 2024; Kaletsky *et al*, 2024). The spontaneous creation of so-called epi-alleles through RNAi-related mechanisms has also been described, albeit that these were not found to be long-lived (Wilson *et al*, 2023). In order to firmly test the possibilities and limitations of RNAi inheritance, a strong framework of the responsible molecular mechanisms is required. Unfortunately, at the molecular level the RNAi inheritance process is not well understood. A number of factors have been implicated in the process, including for instance RNA helicases (Dai *et al*, 2022; Wan *et al*, 2018; Ishidate *et al*, 2018), Argonaute proteins (Buckley *et al*, 2012b; Schreier *et al*, 2022; Wan *et al*, 2018; Conine *et al*, 2013; Liu *et al*, 2023a) and potential scaffold proteins of germ granules (Wan *et al*, 2021; Placentino *et al*, 2021; Schreier *et al*, 2022; Ouyang *et al*, 2019; Lev *et al*, 2019). However, these have not yet been woven into a coherent framework of activities that explain what is happening at the molecular level. An important aspect to note is that these studies made use of self-fertilizating hermaphrodites to study inheritance. While this is a valid approach, it does not allow conclusions on whether a given factor acts in the parents or in the offspring or both. As this is an important first step towards dissecting inheritance mechanisms, we developed an assay in which inheritance through male and female germlines can be seperately studied, by using crosses between males and hermaphrodites. We then used this assay to study factors that have been implicated in RNAi inheritance: HRDE-1, WAGO-1, WAGO-3, WAGO-4, ZNFX-1, PEI-1 and PEI-2. Our results lead us to propose a pathway for RNAi inheritance in *C. elegans*.

## RESULTS

### Exogenous RNAi effects are heritable via both oocyte and sperm

Although RNAi through the feeding of dsRNA-expressing bacteria has been shown to work robustly for *C. elegans* hermaphrodites, its effectiveness is thought to be very low in males (Bezler *et al*, 2019b)(Bezler *et al*, 2019a). Nevertheless, to further dissect male, female and zygotic components of the RNAi inheritance pathway, we aimed to develop a feeding-based RNAi protocol for males and hermaphrodites. We generated a strain that specifically expressed a *gfp::histone-H4* transgene as an RNAi target throughout male and female germline development, and in mature gametes. The transgene contained a *his-67* promoter, driving a *gfp* coding sequence containing three introns fused to *his-67* followed by the 3’UTR from the *tbb-2* gene. This strain also contained the *him-5(e1490)* mutation to increase the frequency of males in the culture and expressed an endogenously tagged PGL-1::mTagRFP-T fusion protein. The latter served for automated germline detection during image processing and normalization of GFP::H4 fluorescence. We treated animals for one generation (embryo until adulthood) with either *control* RNAi or *gfp* RNAi via feeding, and calculated the relative fluorescence intensity (RFI; see Methods) of GFP::H4 in adult hermaphrodites and adult males using microscopy (Figure 1A). We found that males were as sensitive to the *gfp* RNAi treatment as hermaphrodites, given that no GFP::H4 signal could be detected with either fluorescence-based or immuno-based approaches (Figure 1B-D).

**Figure 1.**
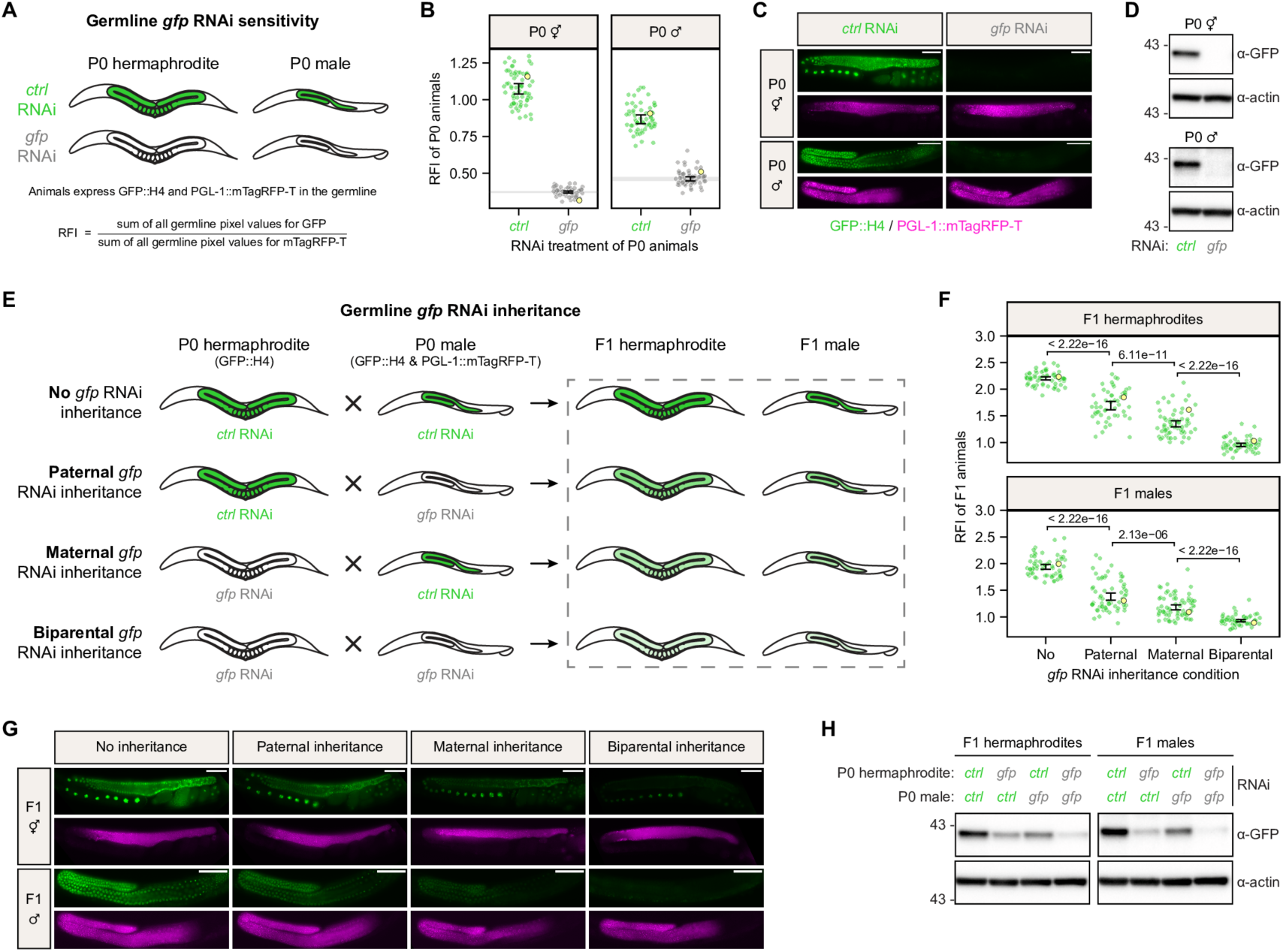
Exogenous RNAi effects are heritable via both oocyte and sperm. A, Schematic representation summarizing the *gfp* RNAi effect on P0 hermaphrodites and P0 males expressing GFP::H4 and PGL-1::mTagRFP-T in the germline. Germline color illustrates GFP::H4 expression: green – expressed, white – silenced. B, Relative fluorescence intensity (RFI) of GFP::H4 in P0 hermaphrodites and P0 males treated with either *control* RNAi (green) or *gfp* RNAi (grey). Each dot represents an individual animal, with yellow dots referring to representative micrographs shown in (C). 95 % confidence intervals of the median are shown as black error bars for all samples, and as grey lines for *gfp* RNAi treatments. Sample size = ∼ 60 animals per condition. C, Widefield fluorescence micrographs of representative P0 hermaphrodites and P0 males treated with either *control* RNAi or *gfp* RNAi, as indicated in (B). GFP::H4 and PGL-1::mTagRFP-T appear in green and magenta, respectively. Scale bars: 50 µm. D, Representative Western blots comparing levels of GFP::H4 with respect to β-Actin loading control in P0 hermaphrodites and P0 males treated with either *control* RNAi or *gfp* RNAi. Sample size = 50 animals per condition. E, Crossing schemes summarizing the effect of *gfp* RNAi inheritance via male and/or female gametes on germline GFP::H4 expression in F1 animals. Germline color illustrates GFP::H4 expression: green – expressed, shades of light green – expressed at lower levels, white – silenced. F, Relative fluorescence intensity (RFI) of GFP::H4 in F1 hermaphrodites and F1 males after *gfp* RNAi inheritance via male and/or female gametes. Each dot represents an individual animal, with yellow dots referring to representative micrographs shown in (G). 95 % confidence intervals of the mean are shown as black error bars. The p-values are from t-tests for linear contrasts in Gaussian models and have been corrected for multiple testing per sex. Sample size = ∼ 60 F1 animals from 5 P0 founders per condition. G, Widefield fluorescence micrographs of representative F1 hermaphrodites and F1 males after *gfp* RNAi inheritance via male and/or female gametes, as indicated in (F). GFP::H4 and PGL-1::mTagRFP-T appear in green and magenta, respectively. Scale bars: 50 µm. H, Representative Western blots comparing levels of GFP::H4 with respect to β-Actin loading control in F1 hermaphrodites and F1 males after *gfp* RNAi inheritance via male and/or female gametes. Sample size = 50 F1 animals from 5 P0 founders per condition.

Next, we set up crosses between differently treated animals to test and quantify the individual contribution of oocytes and sperm to *gfp* RNAi inheritance. In order to identify progeny produced by allogamy, only males expressed both GFP::H4 and PGL-1::mTagRFP-T (hereafter referred to as P0 males), while hermaphrodites only expressed GFP::H4 and were wild-type for *him-5* and *pgl-1* (hereafter referred to as P0 hermaphrodites). Thus, only F1 animals that were produced by mating expressed PGL-1::mTagRFP-T. Using these two strains, we crossed *control* or *gfp* RNAi-treated P0 hermaphrodites with RNAi-treated P0 males in a way that either i) no parent, ii) only P0 males, iii) only P0 hermaphrodites or iv) both parents were treated with *gfp* RNAi (Figure 1E). We found that exogenous RNAi effects were faithfully inherited via both oocytes and sperm (Figure 1F-H). Furthermore, our data shows that *gfp* RNAi inheritance via the oocyte was slightly more effective than via sperm, but less effective compared to an inheritance via both gametes. These findings were independent of the sex of the progeny, as they were observed for both F1 hermaphrodites and F1 males (Figure 1F-H). We note that the *gfp* RNAi effects can also be transgenerationally inherited by following autogamy of P0 hermaphrodites, just as described in previous studies (Figure S1A-C) (Buckley *et al*, 2012b)(Buckley *et al*, 2012a).

### WAGO-4 and HRDE-1 affect germline *gfp* RNAi establishment in hermaphrodites

Having established this exogenous RNAi inheritance assay that allows us to individually investigate maternal or paternal contributions, we sought to examine the requirement of proteins that were previously shown to act in RNAi inheritance: WAGO-1, WAGO-3, WAGO-4, HRDE-1 and ZNFX-1 (Conine *et al*, 2010; Schreier *et al*, 2022; Wan *et al*, 2018; Buckley *et al*, 2012b; Liu *et al*, 2023b). Firstly, we investigated their role in maternal *gfp* RNAi inheritance and started by assessing the effect of each mutant on germline *gfp* RNAi sensitivity in adult hermaphrodites (Figure 2A-K). Along with their respective wild-type controls, we quantified the fluorescence intensity of GFP::H4 in mutant animals that were treated with either *control* RNAi or *gfp* RNAi. We found that *wago-1*, *wago-3* and *znfx-1* mutants did not show any defects in germline GFP::H4 silencing (Figure 2B-D,G-I). In contrast, germline-wide RNAi defects in *wago-4* mutants became occasionally apparent after a few mutant generations and further intensified over time (Figure 2E,J, Figure S2A-F). *hrde-1* mutants also showed impaired RNAi sensitivity, as they consistently failed to silence GFP::H4 in the distal germline (Figure 2F,K). These data highlight functions for WAGO-4 and HRDE-1 in the establishment of germline RNAi. While WAGO-4 is seemingly required to maintain germline RNAi sensitivity over generations, HRDE-1 is specifically required for exogenous RNAi-mediated gene silencing in the distal germline. We also note that the basal expression of the transgene appeared to be affected by specific mutations. This aspect will be covered in more detail later.

**Figure 2.**
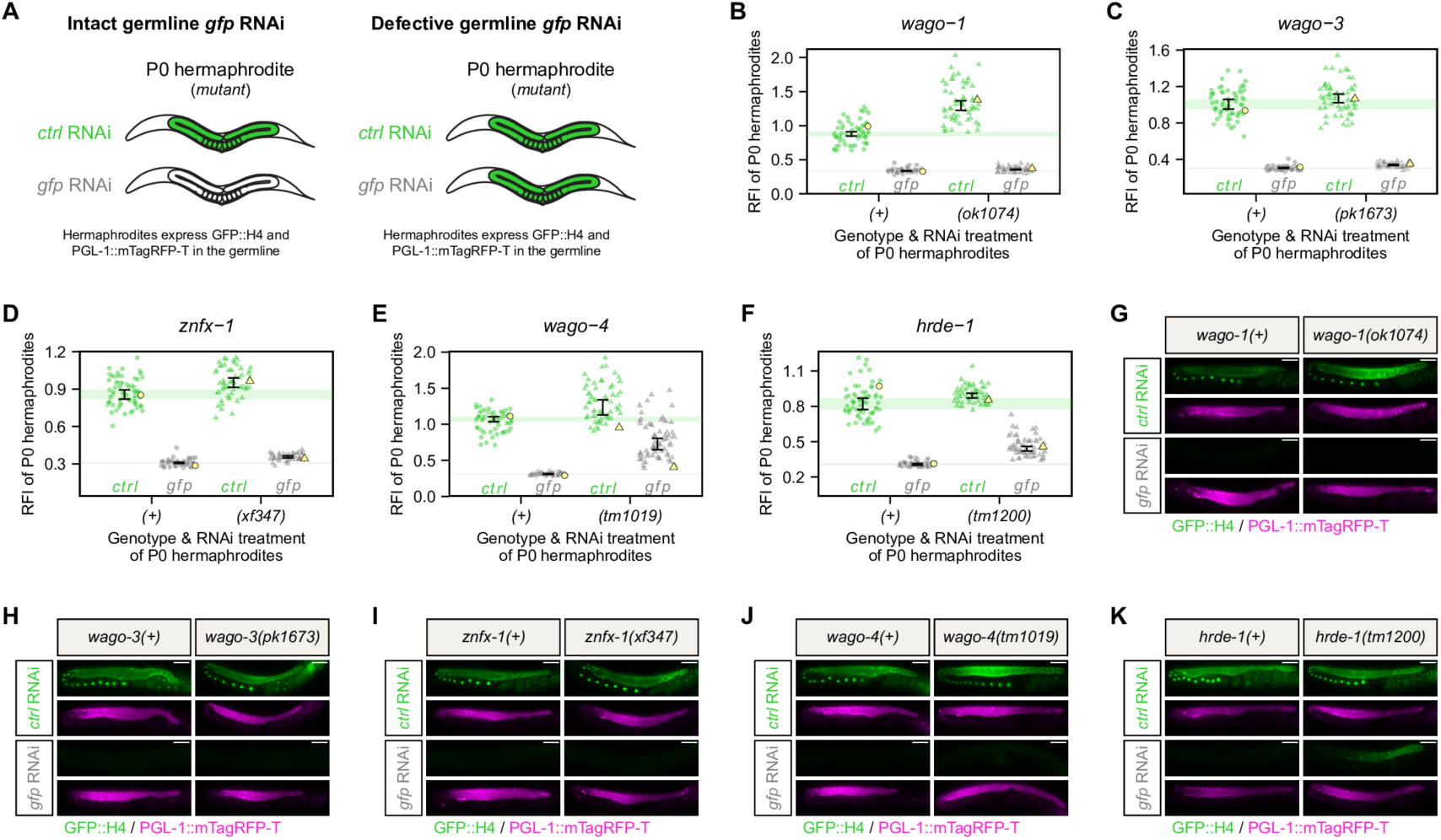
WAGO-4 and HRDE-1 are required for germline RNAi establishment in hermaphrodites. A, Schematic representations summarizing the *gfp* RNAi effect on mutant P0 hermaphrodites expressing GFP::H4 and PGL-1::mTagRFP-T in the germline. Germline color illustrates GFP::H4 expression: green – expressed, white – silenced. B-F, Relative fluorescence intensity (RFI) of GFP::H4 in P0 hermaphrodites treated with either *control* RNAi (green) or *gfp* RNAi (grey). P0 hermaphrodites were either wild-type (circle) or mutant (triangle) for *wago-1* (B), *wago-3* (C), *znfx-1* (D), *wago-4* (E) or *hrde-1* (F). Each dot represents an individual animal, with yellow dots referring to representative micrographs shown in (G-K). 95 % confidence intervals of the median are shown as black error bars for all samples as well as green (*control* RNAi) and grey (*gfp* RNAi) lines for wild-type conditions. Sample size = ∼ 60 animals per condition. G-K, Widefield fluorescence micrographs of representative P0 hermaphrodites treated with either *control* RNAi or *gfp* RNAi, as indicated in (B-F). P0 hermaphrodites were either wild-type or mutant for *wago-1* (G), *wago-3* (H), *znfx-1* (I), *wago-4* (J) or *hrde-1* (K). GFP::H4 and PGL-1::mTagRFP-T appear in green and magenta, respectively. Scale bars: 50 µm.

### Maternal WAGO-3 and WAGO-4, but not HRDE-1 and ZNFX-1 are required for RNAi inheritance via the oocyte

As all investigated mutants succeeded to silence GFP::H4 during oogenesis, we continued by examining their effect on maternal *gfp* RNAi inheritance (Figure 3A). To do this, we crossed wild-type and mutant P0 hermaphrodites treated with either *control* RNAi or *gfp* RNAi with P0 males that were wild-type for the respective genes and treated with *control* RNAi, taking care that RNAi-triggering bacteria were transferred as little as possible (see Methods). Both males and hermaphrodites were homozygous for the GFP::H4 transgene. F1 animals of all four crosses were grown until adulthood and GFP::H4 fluorescence was quantified using microscopy. We then calculated the relative GFP::H4 fluorescence reductions and compared them between F1 animals sired by wild-type and mutant P0 hermaphrodites. In Figures 3B and 3C we present a statistical summary of the data, as described in more detail in the Methods section. The data behind these plots is presented in the other panels (Figure 3-M). Surprisingly, we found that maternal loss of ZNFX-1 caused the maternally inherited *gfp* RNAi effect to be significantly stronger, as we measured a greater GFP::H4 fluorescence reduction in both F1 hermaphrodites and F1 males (Figure 3B,D,I). Also contrary to our expectations, we found that *hrde-1* mutant P0 hermaphrodites still passed on *gfp* RNAi, to both F1 hermpaphrodites and males, and also *wago-1* mutants behaved like wild-type (Figure 3B,F,K). Finally, F1 animals sired by *wago-3* or *wago-4* mutant hermaphrodites revealed clear RNAi inheritance defects, independent of the filial sex (Figure 3B,C,G-H,L-M). Given the observation that *wago-4* mutant hermaphrodites become resistant to germline RNAi over generations (Figure S2A-F), we also tested whether the duration of *wago-4* mutant homozygosity affected maternal *gfp* RNAi inheritance. To achieve this, we compared P0 hermaphrodites carrying the *wago-4* mutation for either four or eight generations for their ability to inherit *gfp* RNAi effects. We found that the duration of *wago-4* mutant homozygosity did mostly not affect RNAi inheritance to the F1 (Figure S2G-J). Inheritance to F1 males was marginally better after four generations than after eight, but this effect may be driven by the increased GFP::H4 expression in the control strain, particularly after four generations (Figure S2G-J). Independently of the duration of mutant homozygosity and filial sex, we noticed that the maternal inheritance defects were not complete and some *gfp* RNAi effects were still transferred (Figure S2K). We conclude that maternal inheritance of *gfp* RNAi requires WAGO-3 and WAGO-4, and is independent of HRDE-1. Furthermore, our data show that maternal ZNFX-1 restricts maternal inheritance of RNAi.

**Figure 3.**
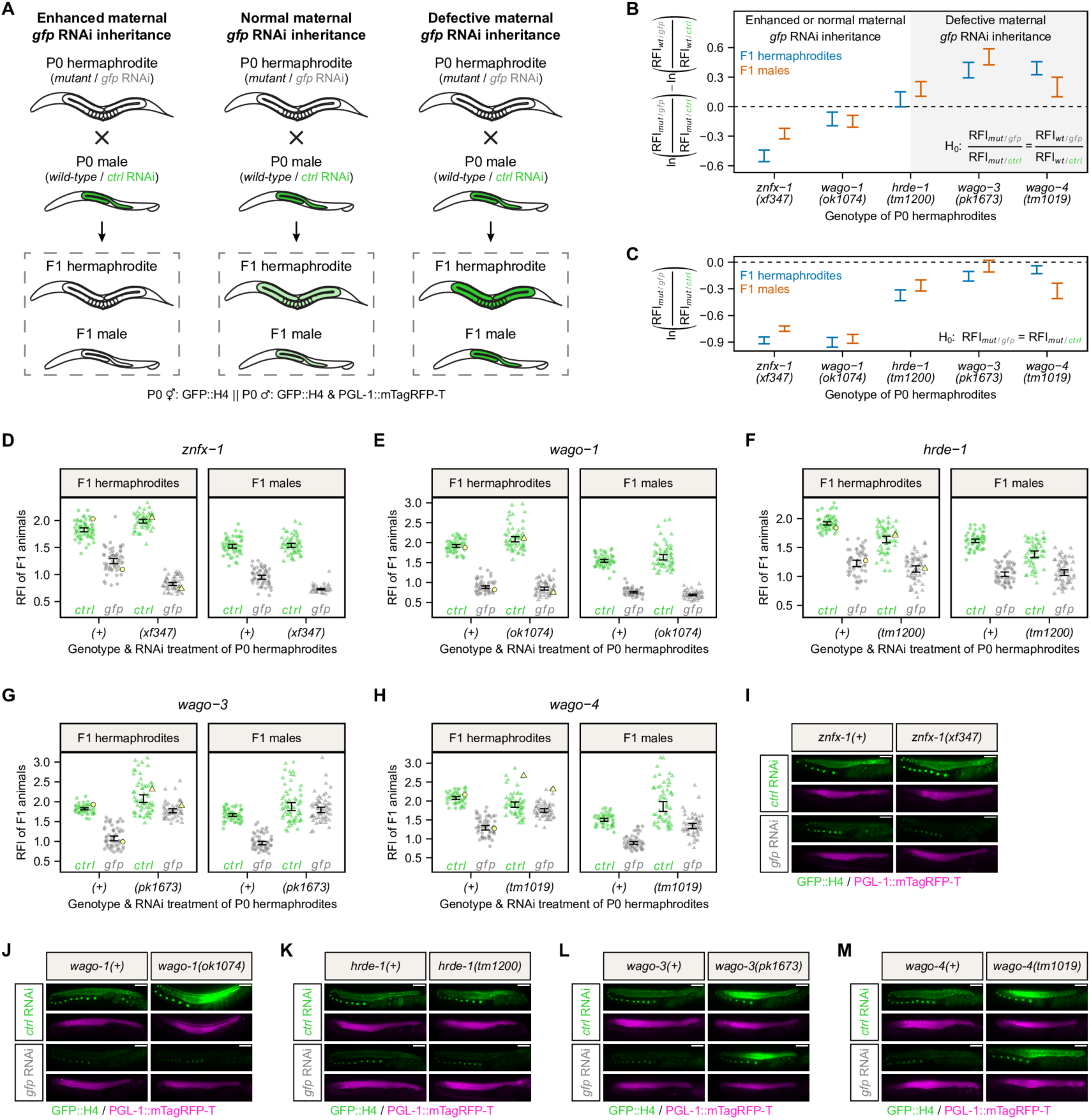
Maternal WAGO-3 and WAGO-4 are required for RNAi inheritance via the oocyte. A, Crossing schemes summarizing how mutations in P0 hermaphrodites affect germline GFP::H4 expression in F1 animals after maternal *gfp* RNAi inheritance. Germline color illustrates GFP::H4 expression: green – expressed, light green – expressed at lower level, white – silenced. B, Comparison of the relative GFP::H4 fluorescence reduction after maternal *gfp* RNAi inheritance between F1 animals sired by wild-type and mutant P0 hermaphrodites for indicated genes. The plot summarizes the Gaussian models fitted in (D-H) and depicts 95 % confidence intervals (CIs) of differences of log fold changes. The null hypothesis (H0) expresses equality of relative GFP::H4 fluorescence reduction between wild-type and mutant condition, meaning that the mutation does not cause an enhanced or defective maternal *gfp* RNAi inheritance. A 95 % CI not including zero is equivalent to a rejection of the null hypothesis at the 5% significance level and indicates that mutations caused either enhanced (95 % CI < 0) or defective (95 % CI > 0) maternal *gfp* RNAi inheritance. Color of CIs indicates sex of F1 animals: blue – hermaphrodite, red – male. C, Comparison of the relative GFP::H4 fluorescence intensity after maternal RNAi inheritance between F1 animals sired by mutant P0 hermaphrodites treated with either *control* RNAi or *gfp* RNAi. The plot summarizes the Gaussian models fitted in (D-H) and depicts 95 % confidence intervals (CIs) of RFI log fold changes. The null hypothesis (H0) expresses equality of relative GFP::H4 fluorescence intensity between *control* RNAi and *gfp* RNAi treatments, meaning that the mutation causes a completely defective maternal *gfp* RNAi inheritance. A 95 % CI not including zero is equivalent to a rejection of the null hypothesis at the 5% significance level and indicates that mutations do not cause a completely defective maternal *gfp* RNAi inheritance. Color of CIs indicates sex of F1 animals: blue – hermaphrodite, red – male. D-H, Relative fluorescence intensity (RFI) of GFP::H4 in F1 hermaphrodites and F1 males after maternal RNAi inheritance. P0 hermaphrodites were treated with either *control* RNAi (green) or *gfp* RNAi (grey), and were either wild-type (circle) or mutant (triangle) for *znfx-1* (D), *wago-1* (E), *hrde-1* (F), *wago-3* (G) or *wago-4* (H). P0 males were always wild-type for these genes and treated with *control* RNAi. Each dot represents an individual animal, with yellow dots referring to representative micrographs shown in (I-M). 95 % confidence intervals of the mean are shown as black error bars. Sample size = ∼ 60 F1 animals from 5 P0 founders per condition. I-M, Widefield fluorescence micrographs of representative F1 hermaphrodites after maternal RNAi inheritance, as indicated in (D-H). Indicated genotypes and RNAi treatments refer to P0 hermaphrodites, which were either wild-type or mutant for *znfx-1* (I), *wago-1* (J), *hrde-1* (K), *wago-3* (L) or *wago-4* (M). GFP::H4 and PGL-1::mTagRFP-T appear in green and magenta, respectively. Scale bars: 50 µm.

### WAGO-4 and HRDE-1 are required for germline *gfp* RNAi in males

We proceeded by investigating the paternal contribution of exogenous RNAi inheritance, for which we also included the PEI granule-localizing proteins PEI-1 and PEI-2 (Schreier *et al*, 2022) in our analyses. Firstly, we assessed the effect of each mutant on germline *gfp* RNAi sensitivity in adult males (Figure 4A-H). Along with their respective wild-type controls, we quantified the fluorescence intensity of GFP::H4 in mutant males that were treated with either *control* RNAi or *gfp* RNAi. We found that *wago-1*, *wago-3*, *znfx-1*, *pei-1* and *pei-2* mutants did not show any defects in germline GFP::H4 silencing in males (Figure 4B-F,I-M). In contrast, *wago-4* and *hrde-1* mutant males showed profound germline RNAi defects, as GFP::H4 fluorescence was still detectable and close to wild-type levels throughout the male germline after *gfp* RNAi treatment (Figure 4G-H,N-O). We conclude that WAGO-4 and HRDE-1 are crucial for the establishment of germline RNAi in males.

**Figure 4.**
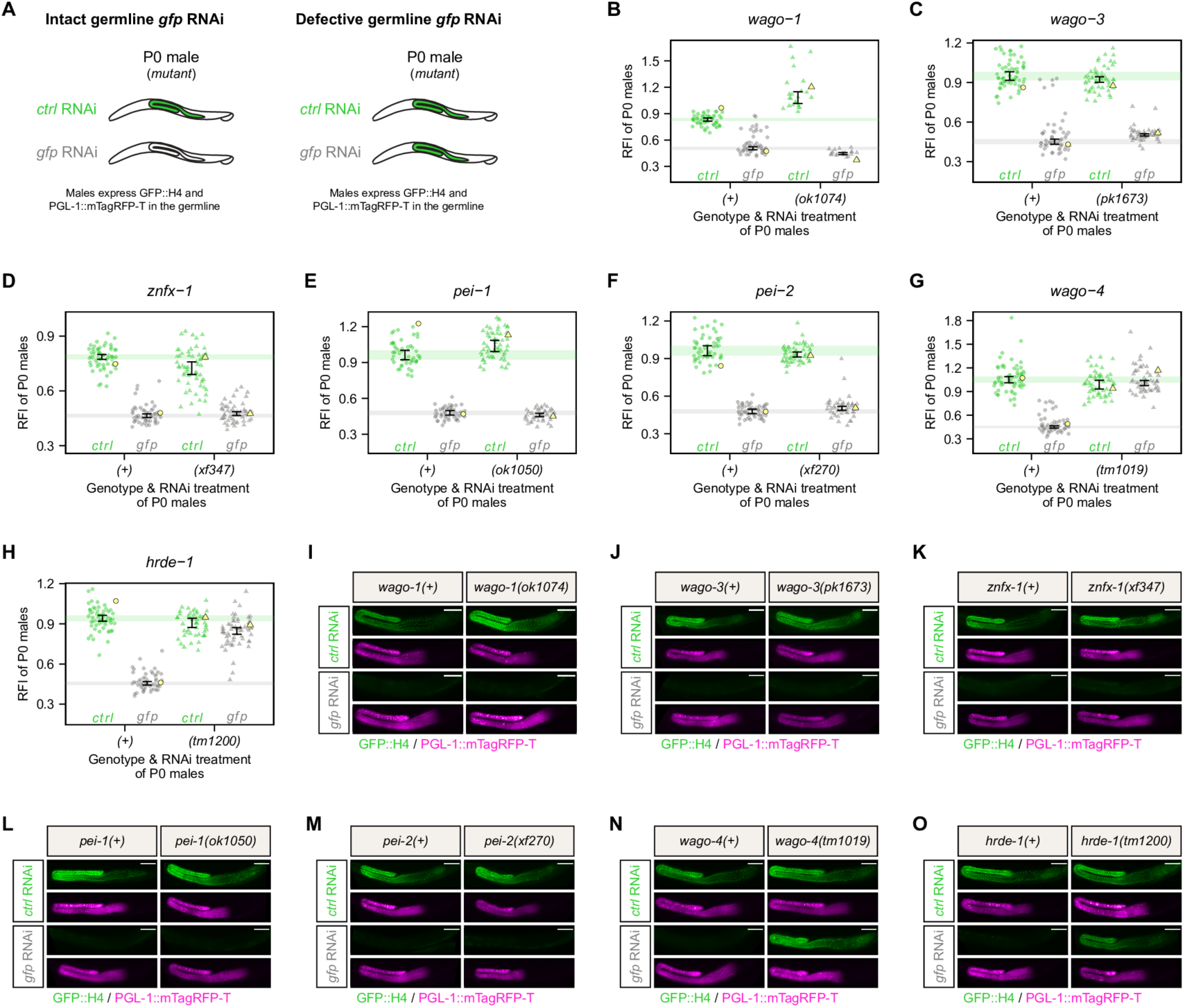
WAGO-4 and HRDE-1 are required for germline RNAi establishment in males. A, Schematic representations summarizing the *gfp* RNAi effect on mutant P0 males expressing GFP::H4 and PGL-1::mTagRFP-T in the germline. Germline color illustrates GFP::H4 expression: green – expressed, white – silenced. B-H, Relative fluorescence intensity (RFI) for GFP::H4 in P0 males treated with either *control* RNAi (green) or *gfp* RNAi (grey). P0 males were either wild-type (circle) or mutant (triangle) for *wago-1* (B), *wago-3* (C), *znfx-1* (D), *pei-1* (E), *pei-2* (F), *wago-4* (G) or *hrde-1* (H). Each dot represents an individual animal, with yellow dots referring to representative micrographs shown in (I-O). 95 % confidence intervals of the median are shown as black error bars for all samples as well as green (*control* RNAi) and grey (*gfp* RNAi) lines for wild-type conditions. Sample size = ∼ 60 animals per condition (exception: ∼ 23 *wago-1(ok1074)* animals). I-O, Widefield fluorescence micrographs of representative P0 males treated with either *control* RNAi or *gfp* RNAi, as indicated in (B-H). P0 males were either wild-type or mutant for *wago-1* (I), *wago-3* (J), *znfx-1* (K), *pei-1* (L), *pei-2* (M), *wago-4* (N) or *hrde-1* (O). GFP::H4 and PGL-1::mTagRFP-T appear in green and magenta, respectively. Scale bars: 50 µm.

### Paternal WAGO-3 and PEI-1 are required for RNAi inheritance via sperm

Next, we investigated the effect of each mutant on paternal *gfp* RNAi inheritance (Figure 5A). We again set up four crosses per investigated gene, where either wild-type or mutant P0 males treated with either *control* RNAi or *gfp* RNAi were crossed with *control* RNAi-treated P0 hermaphrodites that were wild-type for respective genes. GFP::H4 fluorescence of adult F1 animals was quantified using microscopy, and used to calculate the relative GFP::H4 fluorescence reduction after paternal RNAi inheritance (Figure 5B-J). Consistent with the inability to establish germline RNAi (Figure 4G-H), we found that *wago-4* and *hrde-1* mutant males sired offspring that failed to repress GFP::H4 expression after paternal *gfp* RNAi treatment (Figure 5B-E,K-L). We also found that *wago-1*, *znfx-1* and *pei-2* mutant P0 males still inherited *gfp* RNAi effects as efficiently as their respective wild-type controls (Figure 5B,F-H,M-O). In contrast, F1 animals sired by *wago-3* or *pei-1* mutant P0 males revealed clear defects in paternal *gfp* RNAi inheritance. Their relative GFP::H4 fluorescence reduction was significantly lower compared to wild-type controls, with *wago-3* mutant P0 males eliciting a stronger defect than *pei-1* mutant P0 males (Figure 5B,I-J,P-Q). We conclude that paternal inheritance of *gfp* RNAi requires WAGO-3 and PEI-1.

**Figure 5.**
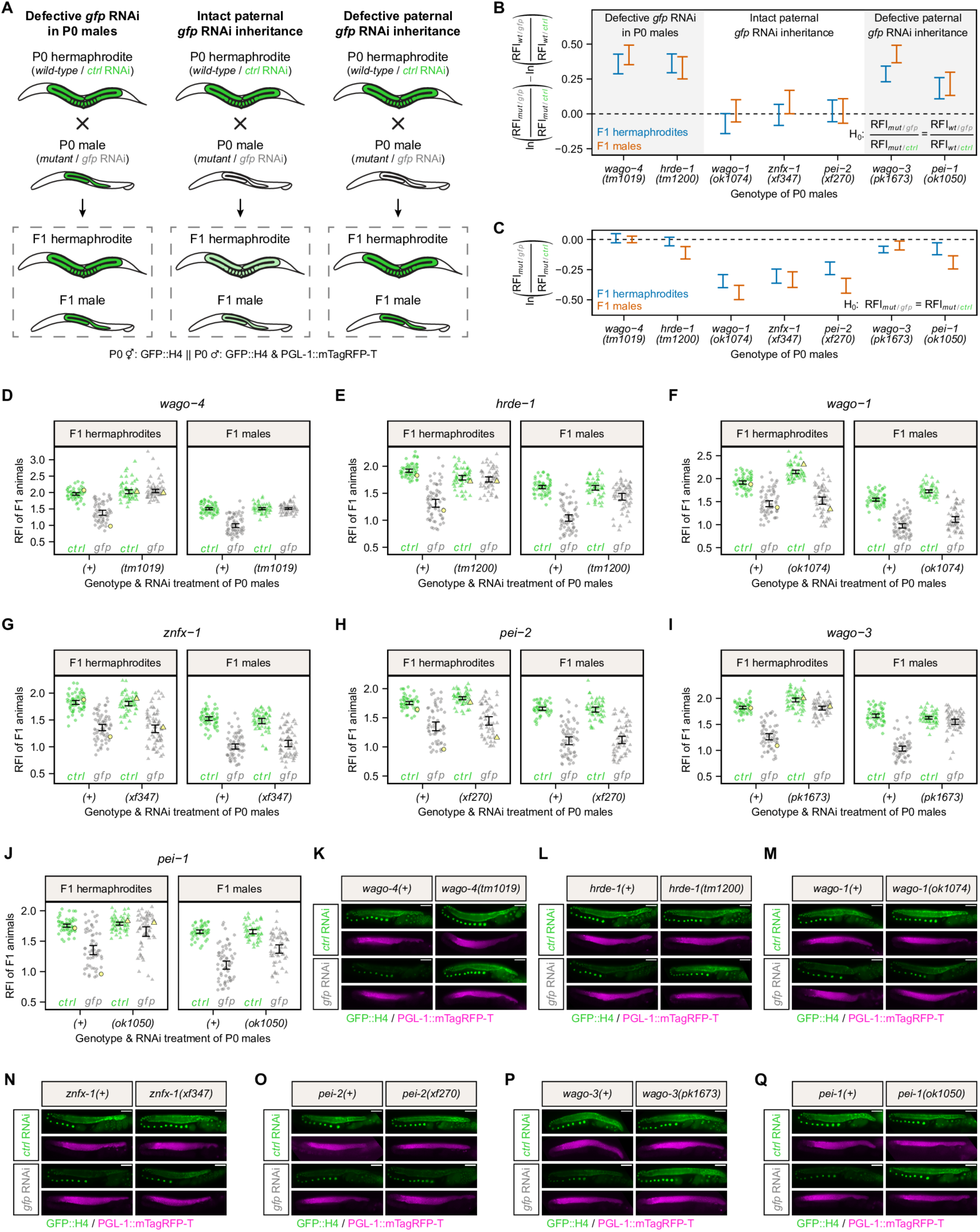
Paternal WAGO-3 and PEI-1 are required for RNAi inheritance via sperm. A, Crossing schemes summarizing how mutations in P0 males affect germline GFP::H4 expression in F1 animals after paternal *gfp* RNAi inheritance. Germline color illustrates GFP::H4 expression: green – expressed, light green – expressed at lower level, white – silenced. B, Comparison of the relative GFP::H4 fluorescence reduction after paternal *gfp* RNAi inheritance between F1 animals sired by wild-type and mutant P0 males for indicated genes. The plot summarizes the Gaussian models fitted in (D-J) and depicts 95 % confidence intervals (CIs) of differences of log fold changes. The null hypothesis (H0) expresses equality of relative GFP::H4 fluorescence reduction between wild-type and mutant condition, meaning that the mutation does not cause a defective paternal *gfp* RNAi inheritance. A 95 % CI not including zero, is equivalent to a rejection of the null hypothesis at the 5% significance level and indicates that mutations caused defective (95 % CI > 0) paternal *gfp* RNAi inheritance. We note that *wago-4* and *hrde-1* mutant P0 males did not inherit the *gfp* RNAi effect due to defective *gfp* RNAi sensitivity (see Figure 4). Color of CIs indicates sex of F1 animals: blue – hermaphrodite, red – male. C, Comparison of the relative GFP::H4 fluorescence intensity after paternal RNAi inheritance between F1 animals sired by mutant P0 hermaphrodites treated with either *control* RNAi or *gfp* RNAi. The plot summarizes the Gaussian models fitted in (D-J) and depicts 95 % confidence intervals (CIs) of RFI log fold changes. The null hypothesis (H0) expresses equality of relative GFP::H4 fluorescence intensity between *control* RNAi and *gfp* RNAi treatments, meaning that the mutation causes a completely defective paternal *gfp* RNAi inheritance. A 95 % CI not including zero is equivalent to the rejection of the null hypothesis at the 5% significance level and indicates mutations did not cause a completely defective paternal *gfp* RNAi inheritance. Color of CIs indicates sex of F1 animals: blue – hermaphrodite, red – male. D-J, Relative fluorescence intensity (RFI) of GFP::H4 in F1 hermaphrodites and F1 males after paternal RNAi inheritance. P0 males were treated with either *control* RNAi (green) or *gfp* RNAi (grey), and were either wild-type (circle) or mutant (triangle) for *wago-4* (D), *hrde-1* (E), *wago-1* (F), *znfx-1* (G), *pei-2* (H), *wago-3* (I) or *pei-1* (J). P0 hermaphrodites were always wild-type for these genes and treated with *control* RNAi. Each dot represents an individual animal, with yellow dots referring to representative micrographs shown in (K-Q). 95 % confidence intervals of the mean are shown as black error bars. Sample size = ∼ 60 F1 animals from 5 P0 founders per condition. K-Q, Widefield fluorescence micrographs of representative F1 hermaphrodites after paternal RNAi inheritance, as indicated in (D-J). Indicated genotypes and RNAi treatments refer to P0 males, which were either wild-type or mutant for *wago-4* (K), *hrde-1* (L), *wago-1* (M), *znfx-1* (N), *pei-2* (O), *wago-3* (P) or *pei-1* (Q). GFP::H4 and PGL-1::mTagRFP-T appear in green and magenta, respectively. Scale bars: 50 µm.

### HRDE-1 and ZNFX-1 are required zygotically to re-establish gene silencing

Our finding that *hrde-1* and *znfx-1* mutant hermaphrodites were able to faithfully inherit *gfp* RNAi contrasts previous studies that reported profound RNAi inheritance defects for these mutants (Buckley *et al*, 2012b; Wan *et al*, 2018). These findings, however, were ascribed by a *gfp* RNAi inheritance assay that scored GFP expression during clonal propagation. As both RNAi-treated animals and non-treated offspring were mutants, defects in either the parental or the filial generation could have caused the defective phenotype. We thus asked whether the strong RNAi inheritance defect described for *hrde-1* and *znfx-1* mutants were based on defective re-establishment of RNAi in F1 animals rather than defective transmission of RNAi effects from the oocyte to the embryo. To test this experimentally, we conducted maternal RNAi inheritance assays, in which we crossed mutant P0 hermaphrodites with either homozygous wild-type or mutant P0 males (Figure 6A). GFP::H4 fluorescence was quantified in adult F1 animals and compared between heterozygous and homozygous mutant offspring. We found that only F1 animals whose parents were both homozygous mutant for *hrde-1* or *znfx-1* showed RNAi inheritance defects (Figure 6B-D); notably, the GFP::H4 signals in these animals were comparable with offspring sired by *control* RNAi-treated P0 hermaphrodites revealing a complete lack of inheritance. Intrigued by these findings, we performed a second RNAi inheritance experiment, in which we aimed to exclude possible effects arising from mutant P0 animals. To achieve this, we set up crosses with heterozygous P0 animals that were treated with *gfp* RNAi. We manually divided the sired F1 animals based on visible GFP expression in the germline (Set 1: GFP negative, Set 2: GFP positive), and used subsets to quantify GFP::H4 fluorescence and occurrence of homozygous mutant F1 animals, respectively (Figure 6E). We found that the group of GFP positive offspring showed a high percentage of homozygous mutants (*hrde-1(tm1200)*: 100 %, *znfx-1(xf347)*: 83.8 %), while GFP negative F1 animals were mostly wild-type or heterozygous for respective genes (*hrde-1(tm1200)*: 1.75 %, *znfx-1(xf347)*: 4.09 %) (Figure 6F-G). We conclude that HRDE-1 and ZNFX-1 are crucial for the re-establishment of gene silencing in the embryo, following the transmission of RNAi effects from the parents.

**Figure 6.**
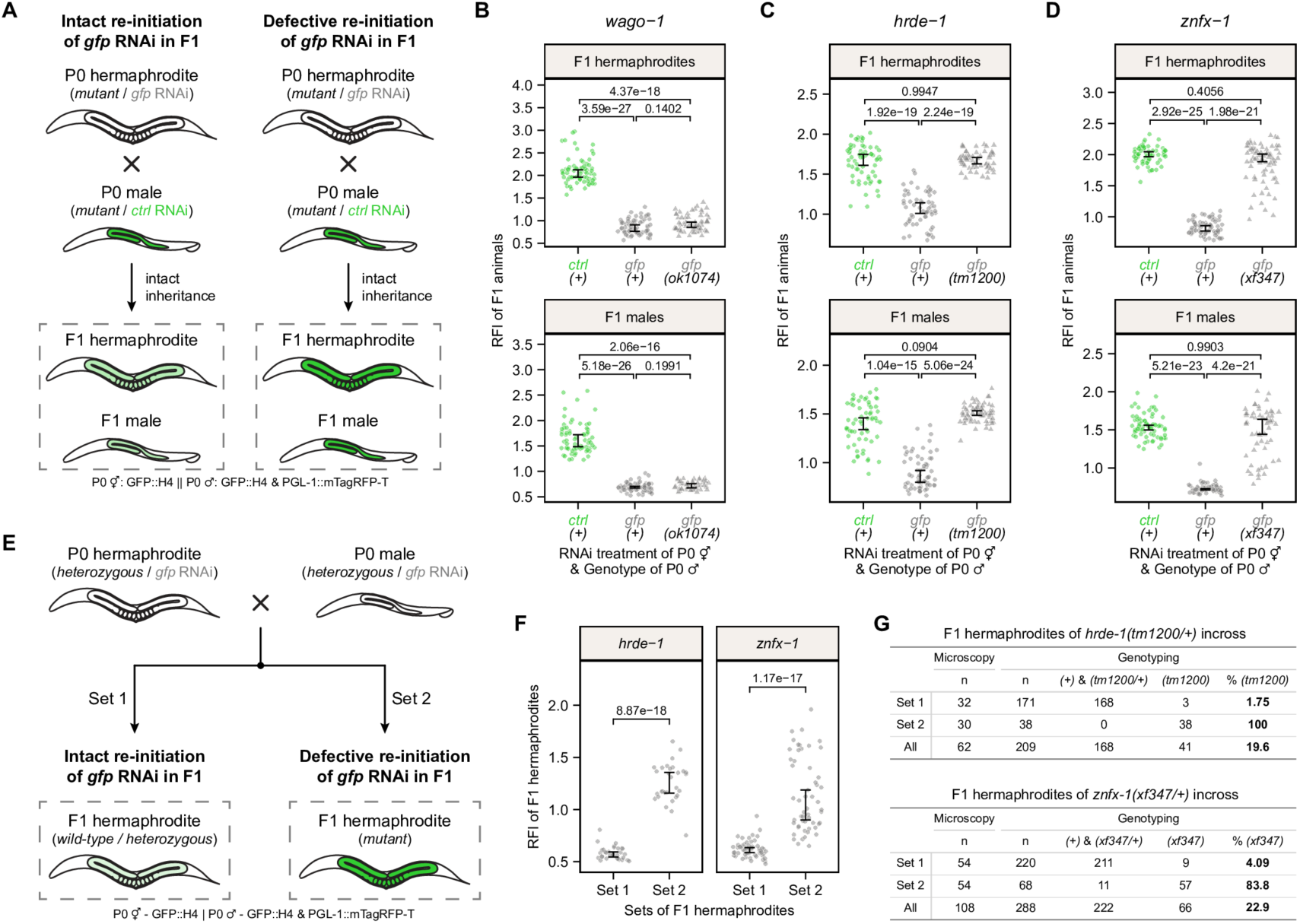
Zygotic HRDE-1 and ZNFX-1 are required for re-initiation of RNAi. A, Crossing schemes summarizing how mutations in F1 animals affect their germline GFP::H4 expression after maternal *gfp* RNAi inheritance. Germline color illustrates GFP::H4 expression: green – expressed, light green – expressed at lower level, white – silenced. B-D, Relative fluorescence intensity (RFI) of GFP::H4 in F1 hermaphrodites and F1 males after maternal RNAi inheritance. P0 hermaphrodites were treated with either *control* RNAi (green) or *gfp* RNAi (grey), and P0 males were either wild-type (circle) or mutant (triangle) for *wago-1* (B), *hrde-1* (C) or *znfx-1* (D). For all crosses, P0 hermaphrodites were mutant for indicated genes and P0 males were treated with *control* RNAi. Each dot represents an individual animal. 95 % confidence intervals of the median are shown as black error bars. Statistically significant differences were determined using two-sided Dunn’s tests. Sample size = ∼ 60 F1 animals from 5 P0 founders per condition. E, Crossing scheme depicting how the genotype of F1 animals affects their germline GFP::H4 expression after biparental *gfp* RNAi inheritance. Germline color illustrates GFP::H4 expression: green – expressed, light green – expressed at lower level, white – silenced. F, Relative fluorescence intensity (RFI) of GFP::H4 in F1 hermaphrodites after biparental *gfp* RNAi inheritance. P0 hermaphrodites and P0 males were treated with *gfp* RNAi and heterozygous for a mutation in either *hrde-1* or *znfx-1*. F1 hermaphrodites were divided in two sets based on visible GFP expression in the germline. Each dot represents an individual animal. 95 % confidence intervals of the median are shown as black error bars. Statistically significant differences were determined using two-sided Wilcoxon rank-sum tests. Sample size = ∼ 31 (*hrde-1*) or 54 (*znfx-1*) F1 animals from 5 P0 founders per condition. G, Genotyping summary and percentage of homozygous mutant F1 animals in each set, as described in (E-F).

### Spatiotemporal regulation of basal GFP::H4 expression in the hermaphroditic germline

We noticed that *wago-1* and *wago-4* mutant P0 hermaphrodites displayed an increased GFP::H4 fluorescence signal in the early/mid pachytene region of the adult germline (Figure 2G,J). In order to describe this unexpected GFP::H4 dysregulation in more detail, we compared mean fluorescence intensities (MFIs) and coefficients of variations (CVs) of GFP::H4 and PGL-1::mTagRFP-T between wild-type and mutant P0 hermaphrodites treated with *control* RNAi (Figure S3A-C). While the MFIs describe overall changes in fluorescence intensity, CVs (ratio of the standard deviation to the mean) describe the dispersion of fluorescence intensities and hence help to determine local alterations. We mainly found minor MFI and CV changes for both GFP::H4 and PGL-1::mTagRFP-T in *wago-3*, *hrde-1* and *znfx-1* mutants. *wago-1* and *wago-4* mutants, however, showed a tendency for elevated MFIs of GFP:H4, but not for PGL-1::mTagRFP-T (Figure S3B). Both mutants also showed a strong and specific increase in CVs for GFP::H4 (Figure S3C), indicating that they indeed expressed GFP::H4 with greater variation compared to their wild-type controls. Our data indicate that the increased fluorescence intensity in the early/mid pachtene region was specific to GFP::H4 in *wago-1* and *wago-4* mutants. Notably, we also found that this altered GFP::H4 pattern is maternally inherited to heterozygous hermaphroditic offspring (Figure 3J,M), but not male offspring (Figure S3D,E). Interestingly, similar increased expression was detected in offspring sired by *wago-3* mutant hermaphrodites and wild-type males, even though this was not detected in the parental generation (Figure 2H, Figure 3L, Figure S3D-E). Finally, we also asked whether these increases in transgene expression were specific to the maternal germline and performed the same analyses with P0 males (Figure S4A-E). This only revealed slight increases in MFIs and CVs for GFP::H4 in *wago-1* mutant males (Figure S4B-C). We conclude that *wago-1* and *wago-4* affect the basal expression of the GFP::H4 transgene, indicating that it is under RNAi-related control also in absence of dsRNA targeting GFP. These effects can be inherited, which is affected by *wago-3*.

## DISCUSSION

We developed an exogenous RNAi inheritance assay that enables the specific and unbiased investigation of the maternal, paternal and filial contribution to RNAi inheritance via the combination of mating experiments and fluorescence quantification of a transgenic GFP::H4 reporter. The assay also allows the simultaneous examination of germline RNAi sensitivity across the various regions of the male and hermaphroditic germlines, including mature gametes. We used this assay to further characterize established RNAi mutants, and successfully dissected formerly reported inheritance defects by attributing specific roles during: i) different germ cell stages in the adult, ii) sperm-mediated transmission of RNAi effects, iii) oocyte-mediated transmission of RNAi effects, and iv) re-establishment of silencing in offspring of RNAi-treated animals (Figure 7). The impact of our results on these four aspects will be discussed separately below.

**Figure 7.**
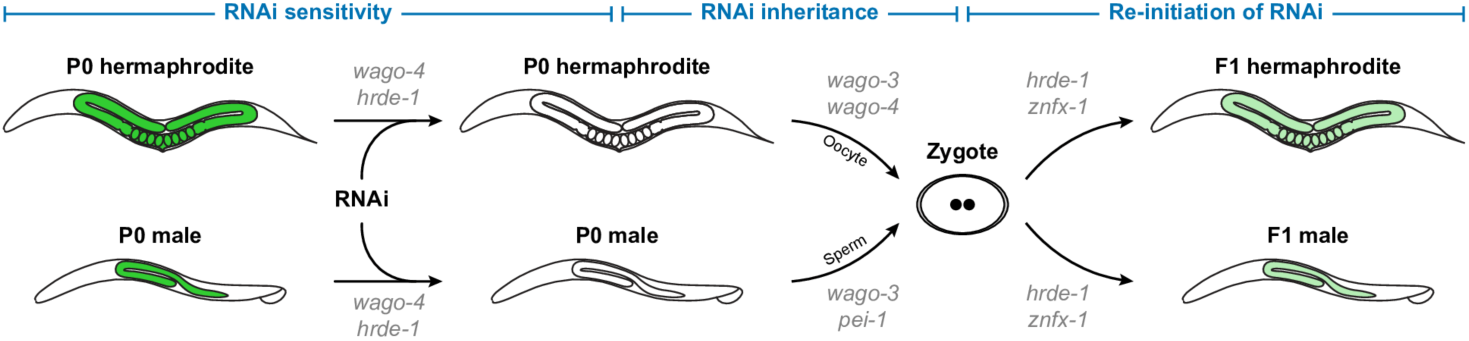
Working model of the three major steps governing exogenous RNAi inheritance. Schematic representation summarizing the three major steps during exogenous RNAi inheritance and their required factors. 1) RNAi sensitivity: Upon exposure to double-stranded RNA, WAGO-4 and HRDE-1 are individually required to mediate gene silencing of the germline-expressed target transcript in both P0 hermaphrodites and P0 males. Interestingly, their impact differs between both sexes. While lack of each protein causes global and immediate germline RNAi sensitivity defects in P0 males, their impact in P0 hermaphrodites is less comprehensive. Here, HRDE-1 is seemingly required for gene silencing in meiotic germ cells, while germline-wide defects in *wago-4* mutants become only apparent after a couple of mutant generations. 2) RNAi inheritance: After successful gene silencing in the germline, WAGO-3 and WAGO-4 are individually required for faithful transmission of RNAi effects via the oocyte to the embryo. Notably, WAGO-3 is also required for RNAi inheritance via the sperm. This paternal contribution seems to be mainly driven by PEI granules, as only *wago-3* and *pei-1* mutants showed defects in paternal RNAi inheritance. 3) Re-initiation of RNAi: Upon faithful transmission of RNAi effects from the gamete(s) to the embryo, HRDE-1 and ZNFX-1 are individually required for the re-initiation of RNAi in both hermaphroditic and male offspring.

### The adult germline

#### RNAi

We detected roles for HRDE-1 and WAGO-4 in the establishment of RNAi in the adult germline, while the other tested factors, including WAGO-1 and WAGO-3, were not required at any stage. HRDE-1 is required for RNAi establishment in the distal region of the hermphroditic germline, as also observed before (Ouyang *et al*, 2022). This implies that in the mitotic and early meiotic stages nuclear RNAi is the main driver of the silencing effect. During later meiotic stages, nuclear RNAi is apparently no longer required, as RNAi triggered silencing effectively in these stages in *hrde-1* mutants. Possibly, this reflects a shift to overall more post-transcriptional gene regulation during the final steps of oogenesis. The involvement of WAGO-4 in RNAi establishment only became apparent after a number of generations (>4) in homozygous mutants. Notably, RNAi establishment in these later generation *wago-4* mutants is defective all over the germline, also in the HRDE-1 dependent zone. This may indicate that WAGO-4 and HRDE-1 functionally interact to set up RNAi in mitotic and early meiotic germ cells. However, given the delay in phenotype establishment the interaction may be rather indirect. For instance, via a destabilization of the overall RNAi network, leading to an overall loss of RNAi effectivity across the whole germline. The endogenous 22G RNAs bound by WAGO-4 are different from those of HRDE-1, further supporting this potential indirect effect of WAGO-4 on HRDE-1 functionality.

In the male germline both HRDE-1 and WAGO-4 are required at all stages. Notably, in *wago-4* mutant males we did not observe the intermediate sensitivity we detected initially in the hermaphrodites, but simply full RNAi resistance. We hypothesize that the male germline may be more sensitive to the effects of WAGO-4 on the RNAi network compared to the female germline.

#### Basal expression of the GFP::H4 transgene

Expression levels of the transgene in absence of RNAi were affected by some of the mutants we tested. In particular WAGO-1 appears to play a major role in this, in both the male and female germline. Strikingly, *wago-1* mutants did not display any defects in RNAi establishment or its inheritance, suggesting that WAGO-1 may be dedicated to endogenous silencing cues. While *wago-4* also affected this aspect, its effects were weaker, and it is feasible that these stem from indirect, RNAi-network-destabilizing effects, similar to what was discussed above. Interestingly, this effect on basal expression was also observed in heterozygous offspring from *wago-3* mutant parents, but not in *wago-3* mutants themselves. How may this be explained? We propose that this effect stems from the sudden introduction of parental small RNAs into an RNAi system that was lacking such parental contributions, as WAGO-3 seems to be the main conveyor of small RNAs between generations (see below). The absence of such parental small RNAs may lead to a re-equilibration of the overall RNAi network and that may be de-stabilized by re-introduction of paternal or maternal small RNAs. This may in turn affect WAGO-1 function. Interestingly, WAGO-1 and WAGO-3 have been shown to interact *in vivo* (Schreier *et al*, 2022; Barucci *et al*, 2020), suggesting both may be present in one larger RNP, which may play a role in the ‘communication’ between these two argonaute proteins. These ideas will need to be tested in future experiments, but they start to rationalize the rather convoluted effects of *wago* mutants that have been observed thus far.

### Sperm-mediated transmission of RNAi

WAGO-3 was implicated in paternal transmission of RNAi. It is not required for establishment, while inheritance via sperm was fully blocked in sperm derived from *wago-3* mutants. Consistent with our previous work on PEI granules that secure WAGO-3 in sperm (Schreier *et al*, 2022), PEI-1 was also found to be required. However, the effect of *pei-1* was less strong. Likely, this reflects the previously described loss of much, but *not all* of the WAGO-3 protein into the residual body during spermatogenesis (Schreier *et al*, 2022). We could not detect a role for the second structural PEI granule component: PEI-2. This may be explained by the fact that in *pei-2* mutants PEI granules are evenly distributed across spematids and residual bodies, leading to even more WAGO-3 remaining in the sperm, and concentrated in PEI granules, compared to *pei-1* mutants (Schreier *et al*, 2022).

Loss of HRDE-1 also blocked inheritance. However, in *hrde-1* mutant males no RNAi was established, making it hard to interpret its role in inheritance. Nevertheless, the following may be deduced from this finding. If we assume that the loading, and or the effects of WAGO-3 and HRDE-1 in response to RNAi would be independent there would be no reason that *hrde-1* mutants would be inheritance-defective. Given that they are, they must be dependent in some manner. For instance, WAGO-3 may be loaded in response to HRDE-1 activity. Indeed, HRDE-1 has been implicated in triggering so-called tertiary 22G RNAs (22G RNAs triggerd by secondary 22G RNAs) (Sapetschnig *et al*, 2015). Alternatively, HRDE-1-driven effects on chromatin may be required to potentiate WAGO-3 activity at later stages of the inheritance process. These and other possibilities will have to be probed in future experiments.

### Oocyte-mediated transmission of RNAi

Inheritance via the oocyte also depends on WAGO-3. Indeed, WAGO-3 is also expressed in the female germline (Schreier *et al*, 2022; Seroussi *et al*, 2023). However, in another assay, named mutator-induced sterility (Mis-assay) (de Albuquerque *et al*, 2015), we could not detect a role for maternal WAGO-3 (Schreier *et al*, 2022). The main difference between the RNAi assay described here and the Mis-assay is that the latter is driven by endogenous small RNAs, while the RNAi assay is driven by exogenous RNAi. Also, the Mis-assay readout likely is a rather convoluted effect, much downstream of the original RNAi-related effects that are being inherited (de Albuquerque *et al*, 2015; Phillips *et al*, 2015). It is therefore feasible that in the Mis-assay loss of WAGO-3 may be compensated by other effects. These may be other Argonaute proteins, or other gene-regulatory events. As the RNAi-assay described here is more direct, it is likely that WAGO-3 is indeed involved in 22G RNA inheritance, both via sperm as well as via oocytes.

Maternal HRDE-1 was not required for inheritance. This differs from the male germline, where HRDE-1 was required. While it does not rule out that HRDE-1 may drive WAGO-3 activity in the female germline, like it does in the male germline, this result does show that WAGO-3 may be loaded in response to another WAGO protein, possibly WAGO-4. WAGO-4 was also found to be involved in oocyte-mediated inheritance. Mutants showed this defect also at early generations, when *wago-4* mutant hermaphrodites did not yet display any RNAi-establishment defects. How the *wago-3* and *wago-4* effects relate to each other is currently unclear, but given that they are apparently not redundant they may act together to drive inheritance. In this light, the effect of maternal ZNFX-1 on inheritance is noteworthy: maternal ZNFX-1 limits inheritance via the oocyte. Interestingly, ZNFX-1 has been described to affect WAGO-4 localization to germ granules (Wan *et al*, 2018), but the functional relevance of this effect is not clear. Based on our data one could hypothesize that maternal ZNFX-1 restricts WAGO-3-mediated 22G RNA inheritance possibly by favouring WAGO-4/WAGO-1-driven pathways (see effects of WAGO-4 and WAGO-1 on basal transgene expression, discussed above). Absence of ZNFX-1 in the mother may thus result in more WAGO-3 being loaded with GFP-targeting 22G RNAs in our set-up. ZNFX-1 has been shown to interact with pUGylated RNAs (Ouyang *et al*, 2022) which serve as the template for 22G RNA biogenesis (Shukla *et al*, 2020) and RdRP (Ishidate *et al*, 2018), which synthesize the 22G RNAs, suggesting that indeed it may act upstream of specific WAGO proteins to direct their loading. Possibly, ZNFX-1 favours the loading of argonaute proteins acting in endogenous pathways over loading of argonaute proteins in response to exogenous dsRNA, such as we provide in our inheritance assays. This would be in line with the previously described balancing role for ZNFX-1 (Ishidate *et al*, 2018). One major hurdle to better understand the effects we describe is that we do not know which WAGO protein(s) is(are) required for establishing RNAi in the late oogenesis stages. None of the tested WAGO proteins are on their own, suggesting that this is driven by a combination of the tested WAGOs and/or by one that we did not test.

### Re-establishment of silencing in offspring of RNAi-treated animals

Zygotic ZNFX-1 was found to be required for the re-establishment of silencing in the embryo, following inheritance. This is opposite to the effect we detected for maternal ZNFX-1 in RNAi re-establishment in the embryo. How can this be reconciled? If we assume, as discussed above, that maternal ZNFX-1 promotes the loading of argonaute proteins in response to endogenous triggers, at the expense of exogenous triggers, the obtained result suggests that inherited WAGO-3 triggers are interpreted as endogenous triggers in the embryo, requiring ZNFX-1 to result in the loading of argonaute that implicates the silencing. As zygotic HRDE-1 was also found to be essential for re-establishment of silencing in the embryo, the argonaute downstream of ZNFX-1 likely is HRDE-1. This places HRDE-1 downstream of WAGO-3, opposite to how WAGO-3 and HRDE-1 are proposed to interact in the adult germline (see discussion above). Effectively, this constitutes a cycle between WAGO-3 and HRDE-1, where HRDE-1 drives the silencing effect and loading of WAGO-3, while WAGO-3 acts to transmit it to a next generation and to again load HRDE-1. ZNFX-1 may act as a gatekeeper in this system to control the entry of 22G RNAs from exogenous triggers, such as ingested dsRNA, versus endogenous triggers. The complete dependence on HRDE-1 in the embryo also shows that re-establishment of RNAi fully depends on nuclear RNAi activity.

## MATERIALS AND METHODS

### *C. elegans* culture and strains

Unless otherwise stated, all worm strains were cultured according to standard laboratory conditions at 20°C on standard Nematode Growth Medium (NGM) plates seeded with *Escherichia coli* OP50 (Brenner, 1974a)(Brenner, 1974b). All strains are in the N2 Bristol background. Every non-‘RFK’ strain was provided by the *Caenorhabditis* Genetics Center (CGC), which is funded by NIH Office of Research Infrastructure Programs (P40 OD010440). A list of all strains used in this study is provided in Supplementary Table 1. Wormbase (Harris *et al*, 2020; Sternberg *et al*, 2024; Davis *et al*, 2022) was extensively used in these studies.

### Mos-1 mediated transgenesis

Mos1-mediated Single Copy Insertion (MosSCI) was used to generate the strain RFK1305; *xfSi255[his-67p::gfp::his-67::tbb-2 3’UTR + Cbr-unc-119(+)] II* (Frøkjær-Jensen *et al*, 2008b, 2012b). This transgene was targeted to the locus *ttTi5605* on chromosome II. The *his-67* promoter and *tbb-2* 3’ UTR sequence were used for ectopic expression in the whole germline. The *gfp* coding sequence including three introns was amplified from pDD282. An amplified DNA fragment containing the sequences of the *his-67* promoter, *gfp* with three introns, *his-67* CDS and *tbb-2* 3’ UTR was cloned into pCFJ350. All plasmids used for microinjection were purified from 4 ml bacterial culture using PureLink™ HiPure Plasmid Miniprep Kit (Art. No. K210011, Invitrogen™), eluted in sterile water and confirmed by enzymatic digestion and sequencing. A plasmid mix containing 50 ng/μl pCFJ601, 10 ng/μl pMA122, 10 ng/μl pGH8, 5 ng/μl pCFJ104, 2.5 ng/μl pCFJ90 and 50 ng/μl of pRFK4197 were injected in both gonads of 20 young adults of the EG6699 strain. The progeny was screened as previously described (Frøkjær-Jensen *et al*, 2012b, 2008b). Successful insertion events were confirmed by Sanger sequencing. The generated strain was out-crossed two times prior to any further cross or analysis. A list of all plasmids used in this study is provided in Supplementary Table 2.

pDD282 was a gift from Bob Goldstein (Addgene plasmid # 66823; http://n2t.net/addgene:66823; RRID:Addgene_66823) (Dickinson *et al*, 2015b)(Dickinson *et al*, 2015a). pCFJ601, pMA122, pGH8, pCFJ90, pCFJ104 and pCFJ350 were gifts from Erik Jorgensen (Addgene plasmid # 34874; http://n2t.net/addgene:34874; RRID:Addgene_34874, Addgene plasmid # 34873; http://n2t.net/addgene:34873; RRID:Addgene_34873, Addgene plasmid # 19359; http://n2t.net/addgene:19359; RRID:Addgene_19359, Addgene plasmid # 19327; http://n2t.net/addgene:19327; RRID:Addgene_19327, Addgene plasmid # 19328; http://n2t.net/addgene:19328; RRID:Addgene_19328, Addgene plasmid # 34866; http://n2t.net/addgene:34866; RRID:Addgene_34866) (Frøkjær-Jensen *et al*, 2012b, 2008b).(Frøkjær-Jensen *et al*, 2008a, 2012a)

### CRISPR/Cas9-mediated genome editing

CRISPR/Cas9-mediated genome editing was used to generate the *znfx-1(xf347)* indel allele (8517 bp deletion + 44 bp insertion; 27 bp downstream of endogenous *znfx-1* start codon) using the *unc-58(e665)* co-conversion strategy (Arribere *et al*, 2014). All protospacer sequences were chosen using CRISPOR (http://crispor.tefor.net) (Haeussler *et al*, 2016) and cloned in either pRFK2411 (plasmid expressing Cas9 + sgRNA(F+E) (Chen *et al*, 2013); derived from pDD162) or pRFK2412 (plasmid expressing only sgRNA(F+E) (Chen *et al*, 2013); derived from pRK2411) via site-directed, ligase-independent mutagenesis (SLIM) (Chiu *et al*, 2004, 2008; Schreier *et al*, 2022). pDD162 (*eft-3p*::*Cas9* + empty sgRNA) was a gift from Bob Goldstein (Addgene plasmid # 47549; http://n2t.net/addgene:47549; RRID:Addgene_47549) (Dickinson *et al*, 2013). SLIM reactions were transformed in Subcloning Efficiency™ DH5α™ Competent Cells (Art. No. 18265017, Invitrogen™) and plated on LB agar plates supplemented with 100 μg/ml ampicillin. All plasmids used for microinjection were purified from 4 ml bacterial culture using PureLink™ HiPure Plasmid Miniprep Kit (Art. No. K210011, Invitrogen™), eluted in sterile water and confirmed by enzymatic digestion and sequencing. A DNA mix containing 50 ng/μl pRFK2588, 30 ng/μl pRFK3358, 30 ng/μl pRFK3359, 30 ng/μl pRFK3360, 30 ng/μl pRFK3361 and 750 mM SJ763 was injected in both gonads of 20 young adults hermaphrodites maintained at 20°C (Schreier *et al*, 2022). Selected F1 *unc* progeny was screened for insertion or deletion by PCR. Successful editing events were confirmed by Sanger sequencing. The generated mutant strain was out-crossed two times prior to any further cross or analysis. A list of all plasmids and protospacer sequences used in this study is provided in Supplementary Table 2 and Supplementary Table 3, respectively.

### RNA interference

RNAi was performed via feeding *Escherichia coli* HT115(DE3) expressing double-stranded RNA (Kamath *et al*, 2003, 2001; Kamath & Ahringer, 2003). The plasmid L4440 served as negative control and was a gift from Andrew Fire (Addgene plasmid # 1654; http://n2t.net/addgene:1654; RRID:Addgene_1654). A *gfp*-specific sequence was amplified from pDD282 and inserted in L4440 to generate pRFK4103, which was used to perform *gfp* RNAi experiments targeting the transgene *xfSi255[his-67p::gfp::his-67::tbb-2 3’UTR + Cbr-unc-119(+)]*. For each RNAi experiment, HT115(DE3) bacteria transformed with either L4440 or pRFK4103 were freshly grown on LB agar plates supplemented with 100 μg/ml ampicillin and 10 μg/ml tetracycline. Following bleaching of mixed-staged worm strains, embryos were directly transferred to RNAi NGM plates (diameter, 90 mm) and grown at 20 °C for one generation. Crosses between L4-staged hermaphrodites and young adult males were set up after three days. Imaging of adult animals was performed after four days.

### RNAi inheritance assays

RFK1305 (*xfSi255*) and RFK1405 (*xfSi255; pgl-1(xf233); him-5(e1490)*) served as standard P0 hermaphrodite strain and standard P0 male strain, respectively. *Pgl-1(xf233[pgl-1::mTagRfp-t])* served as mating control to identify progeny produced by allogamy. A deletion allele of a gene of interest (*goi*) was crossed into both standard P0 strains. The resulting four strains were used to test the effect of the *goi* on specific steps during RNAi inheritance. Each P0 hermaphrodite strain was imaged twice, once in a *pgl-1(+)* and once in a *pgl-1(xf233[pgl-1::mTagRfp-t])* background. While the former was eventually used for mating, the latter was used to analyze fluorescence data according to the established image processing and statistical analysis protocols, which rely on PGL-1::mTagRFP-T. We note that we did not observe any differences between both *pgl-1* genotypes with regard to *gfp* RNAi sensitivity in any P0 hermaphrodite strain. However, we noticed that *wago-1(ok1074); pgl-1(xf233)* double mutants become sterile after three generations.

Paternal *gfp* RNAi inheritance was tested by crossing standard P0 hermaphrodites treated with *control* RNAi (*control* RNAi / *goi(+)*) with four different P0 males of the following conditions: i) *control* RNAi / *goi(+)*, ii) *gfp* RNAi / *goi(+)*, iii) *control* RNAi / *goi(-)*, iv) *gfp* RNAi / *goi(-)*. These four conditions were also used to assess the effect of the *goi* on *gfp* RNAi sensitivity in P0 males.

Maternal *gfp* RNAi inheritance was tested by crossing standard P0 males treated with *control* RNAi (*control* RNAi / *goi(+)*) with four different P0 hermaphrodites of the following conditions: i) *control* RNAi / *goi(+)*, ii) *gfp* RNAi / *goi(+)*, iii) *control* RNAi / *goi(-)*, iv) *gfp* RNAi / *goi(-)*. These four conditions were also used to assess the effect of the *goi* on *gfp* RNAi sensitivity in P0 hermaphrodites.

Re-initiation of *gfp* RNAi in F1 animals was tested by two different approaches. First, we analyzed GFP::H4 expression in homozygous mutant F1 animals (*goi(-)*) after maternal *gfp* RNAi inheritance. Therefore, we performed a maternal *gfp* RNAi inheritance assay, in which we combined conditions iii) and iv) as described above with the following condition: v) P0 hermaphrodites (*gfp* RNAi / *goi(-)*) crossed with P0 males (*control* RNAi / *goi(-)*). Second, we set up a single cross, in which both P0 hermaphrodites and P0 males were heterozygous for the *goi* and treated with *gfp* RNAi (*gfp* RNAi / *goi(+/-)*). F1 animals expressing PGL-1::mTagRFP-T were sorted into two groups based on presence or absence of visible GFP::H4 expression in the germline. Both groups were split into two subgroups, with the first being used for genotyping and the second being used for microscopy.

For each cross, five L4-staged hermaphrodites and ten young adult males grown for three days on RNAi NGM plates were used. To minimize the potential transfer of RNAi-triggering HT115(DE3) bacteria, each animal was hand-picked into a drop of 100 µl M9 buffer, washed thoroughly and subsequently transferred onto standard NGM plates seeded with *Escherichia coli* OP50 (diameter, 35 mm). After one hour, the RNAi effect was confirmed by microscopy, all animals were combined on a standard NGM mating pate (diameter, 35 mm) and grown overnight at 20°C. On the next day, adult males were hand-picked for single-worm-lysis and adult hermaphrodites were singled onto individual standard NGM plates (diameter, 35 mm) and grown at 20°C. We note that we never detected HT115(DE3) colonies on mating plates. After three days, P0 hermaphrodites and P0 males were genotyped for confirmation. On day later, 25 F1 hermaphrodites and 25 F1 males expressing mTagRFP-T in the germline were hand-picked from each plate, while combining all F1 animals of the same sex for subsequent microscopy (75 animals per slide) and Western blot (50 animals per sample).

Transgenerational *gfp* RNAi inheritance was performed using the standard P0 hermaphrodite strain (*xfSi255*). P0 animals were treated with either *control* RNAi or *gfp* RNAi for one generation as described above. After four days of RNAi treatment, F1 embryos were obtained by bleaching and transferred onto standard NGM plates seeded with *Escherichia coli* OP50 (diameter, 90 mm). For each filial generation, ten random hermaphrodites of the post-*control* RNAi condition and ten hermaphrodites with lowest GFP::H4 expression of the post-*gfp* RNAi condition were singled onto fresh NGM plates seeded with *Escherichia coli* OP50 (diameter, 90 mm) to establish the next generation. Animals were grown at 20°C and 75 adult hermaphrodites per sample were used for microscopy. The experiment was stopped after six filial generations.

### Western Blot

Per sample, 50 adults animals, either hermaphrodites or males, were hand-picked in 1x Novex™ NuPAGE™ LDS sample buffer (Art. No. NP0007, Invitrogen™) supplemented with 100 mM DTT and incubated for 30 min at 95°C. Following thorough mixing and centrifugation for 10 min at 21,000 x g, supernatants were transferred into fresh tubes and stored at -20°C until usage. Together with Color Prestained Protein Standard, Broad Range (10-250 kDa, Art. No. P7719S, New England BioLabs®), samples were separated on a Novex™ NuPAGE™ 10 % Bis-Tris Mini-Protein-Gel (Art. No. NP0301, Invitrogen™) in 1x Novex™ NuPAGE™ MOPS SDS Running Buffer (Art. No. NP0001, Invitrogen™) at 50 mA. Afterwards, proteins were transferred on an Immobilon™-P Membran (PVDF, 0.45 μm, Art. No. IPVH00010, Merck Millipore) for 16 h at 15 V using a Mini Trans-Blot® Cell (Art. No. 1703930, Bio-Rad) and 1x NuPAGE™ Transfer Buffer (Art. No NP0006, Invitrogen™) supplemented with 20 % methanol. Following incubation in 1x PBS supplemented with 5 % skim milk and 0.05 % Tween®20 for 1 h, the PVDF membrane was incubated in 1x PBS supplemented with 0.5 % skim milk, 0.05 % Tween®20 and the primary antibody (1:1,000 monoclonal mouse anti-GFP (B-2), Art. No. sc-9996, Santa Cruz Biotechnology® / 1:5,000 monoclonal rabbit β-Actin (D6A8), Art. No. 8457, Cell Signaling Technology®) for 1 h, followed by three washes with 1x PBS supplemented with 0.05 % Tween®20 (hereinafter referred to as 0.05 % PBS-T) for 10 min each, one hour incubation in 0.05 % PBS-T supplemented with the secondary antibody (1:10,000 anti-mouse IgG, HRP-linked antibody, Art. No. 7076, Cell Signaling Technology® / 1:10,000 anti-rabbit IgG, HRP-linked antibody, Art. No. 7074, Cell Signaling Technology®) and three final washes with 0.05 % PBS-T for 10 min each. Chemiluminescence detection was performed using Amersham™ ECL Select™ Western Blotting Detection Reagent (Art. No. RPN2235, GE Healthcare) and a ChemiDoc™ XRS+ System (Art. No. 1708265, Bio-Rad).

### Microscopy

Per slide, 75 adult hermaphrodites or males were hand-picked into a drop of 100 µl M9 buffer, washed thoroughly and individually transferred to a drop of 50 μl M9 buffer supplemented with 40 mM sodium azide on a coverslip. After 20 min, excess buffer was removed and a glass slide containing a freshly made 2 % agarose (w/v) pad was placed on top of the coverslip. Animals were immediately imaged using a THUNDER Imager (Leica) inverted widefield microscope equipped with a 10x/0.32 dry objective for tile scan acquisition or 40x/0.95 dry objective for individual animal acquisition. All images were acquired in 16-bit format.

For hermaphrodites that were not expressing PGL-1::mTagRFP-T (Appendix Figure S1), tile scans were processed as described above with the following changes: i) mean threshold was used to generate binary files, ii) pencil tool was used to manually separate connecting animals.

### Image processing

ImageJ v.1.54f was used to process all tile scans and images. For each tile scan, the red emission channel (detecting PGL-1::mTagRFP-T) was duplicated twice. One duplicate was converted to an 8-bit color image, while the second duplicate was used to generate a binary file using the triangle threshold. The dilate function was applied to binary files showing hermaphrodites. If required, this process was repeated with other tile scans of the same microscopy slide and duplicated worms were removed. Following merge of the 8-bit color image with the binary file, the pencil tool (set color picker to white for both for- and background, set pencil width = 2) was used to manually separate connected germlines from different animals, or germlines from air bubbles, in the binary file using the signal of the 8-bit color image as reference. After splitting channels, the adjusted binary file was used to identify individual germlines using the analyze particles function with a minimum size threshold to exclude speckles and embryos. Using the ROI manager, overlapping germlines were removed and individual gonads of the same animal were combined (for hermaphrodites only). All regions of interest were rechecked, saved and applied to both the green and red emission channels of the original tile scan file(s) to measure the following values: area, mean, standard deviation, modal, median, min, max and integrated density.

### Statistical analyses

#### Ratios of integrated fluorescence densities were analyzed

The GFP::H4 fluorescence signal was of primary interest for the analysis. However, the integrated density (RawIntDen; sum of pixel values) was susceptible to fluctuations in individual body and gonad size and the mean intensity was susceptible to positional effects. We therefore used the ratio of integrated fluorescence densities,

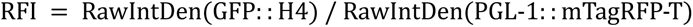

which corrects for both biases simultaneously. Note that a ratio image generated by the microscopy software should not be used, as the fine morphological structure of the GFP and RFP signals is systematically different.

#### F1 data sets were analyzed with Gaussian models in log-scale (*Figures 1*, 3, 5 and S2H-K)

We used Gaussian linear models with categorical predictors for RFI values in logarithmic scale. The normal distribution of the log(RFI) data was checked for each data set using Q-Q plots and Shapiro-Wilk tests. In a few cases the normality assumption was borderline. Due to the large sample size (n > 50), the use of Gaussian models was also unproblematic in these cases. Since the homoscedasticity assumption was violated, we fitted models with inverse variance weights. The large sample size makes this an effective method of correcting for heteroskedasticity. One advantage of the analysis in log scale was that statistical hypothesis tests provided information on relative group differences (log fold changes). In the figures the data are presented in original scale, with the end points of the confidence intervals transformed to original scale. Details of the models are given in the following paragraphs (hereafter ’Gaussian models’). Models were fitted using the ’lm’ function in R 4.3.1 and confidence intervals were calculated using the emmeans v1.8.7 package (Lenth, 2023).

The confidence intervals in Figure 1F are from one-way Gaussian models with the predictor *inheritance type*. They were fitted separately for hermaphrodites and males. The p-values belong to tests of linear contrasts

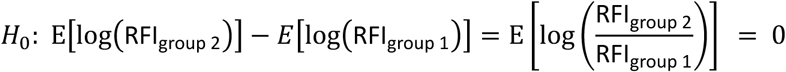

and were corrected for multiple testing (emmeans method ’mvt’, based on multivariate t-distribution).

The data in Figures 3, 5 and S2H-K were analyzed with 2-way Gaussian models with interaction term (predictors *genotype* and *RNAi treatment*). Error bars in panels 3D-H, 5D-J and S2H-I represent 95% confidence intervals for the group means. Panels 3B-C, 5B-C and S2J-K show the results of group comparisons. We present the results of the hypothesis tests as confidence intervals of the interaction parameter

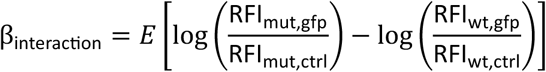

or linear contrasts. This presentation provides more information and is easier to interpret than p-values. Note that if a confidence interval does not contain the value 0, this corresponds to a rejection of the null hypothesis stated in the panel.

#### P0 data sets were analyzed with non-parametric methods (Figures 1B, 2, 4, S1, S2B, S3 and S4)

For data sets where no group comparisons were made or where the assumption of normal distribution was violated, we calculated confidence intervals using the bootstrapping method. In each case, Bias-Corrected and Accelerated Confidence Intervals for the median were calculated using the R-package boot v1.3-28.1 (method ’BCa’, R = 5000) (Canty & Ripley, 2022). The F1 data in Figure 6 was also analyzed with non-parametric model. For group comparisons appropriate non-parametric tests were used. For data sets with 2 groups (Fig. 6F, S3B-E, S4B-E), Wilcoxon rank sum tests were used. For data sets with more than 2 groups (Fig. 6B-D) we used Dunn’s test and corrected for multiple testing with Holm’s method.

## ACKNOWLEDGEMENTS

We thank all members of the Ketting laboratory for critical reading of the manuscript. Joana Costa Pereirinha and Ida Josefine Isolehto are thanked for providing the MosSCI strain expressing GFP::H4 in the whole germline and generating the appropriate *gfp* RNAi clone, respectively. Aaron Noah Ottmann is acknowledged for his contribution to the early stages of this project. We thank the IMB Media Laboratory and Microscopy Core Facility for consumables and equipment. Some strains were provided by the *Caenorhabditis* Genetics Center (CGC), funded by NIH Office of Research Infrastructure Programs (P40 OD010440). This work was funded by the Deutsche Forschungsgemeinschaft (DFG, German Research Foundation) project ID 252386272 to R.F.K..

## AUTHOR CONTRIBUTIONS

J.S. and R.F.K. conceived the study. J.S. designed and executed experiments, performed image processing and statistical analyses. F.K. supervised and performed statistical analyses. R.F.K. supervised the project. J.S. and R.F.K. wrote the manuscript with input from F.K..

## DATA AVAILABILITY

All data needed to evaluate the conclusions in the paper are present in the paper and/or the supplementary materials. Source data are provided with the manuscript. This study includes no data deposited in external repositories.

---

**Appendix Figure S1.**
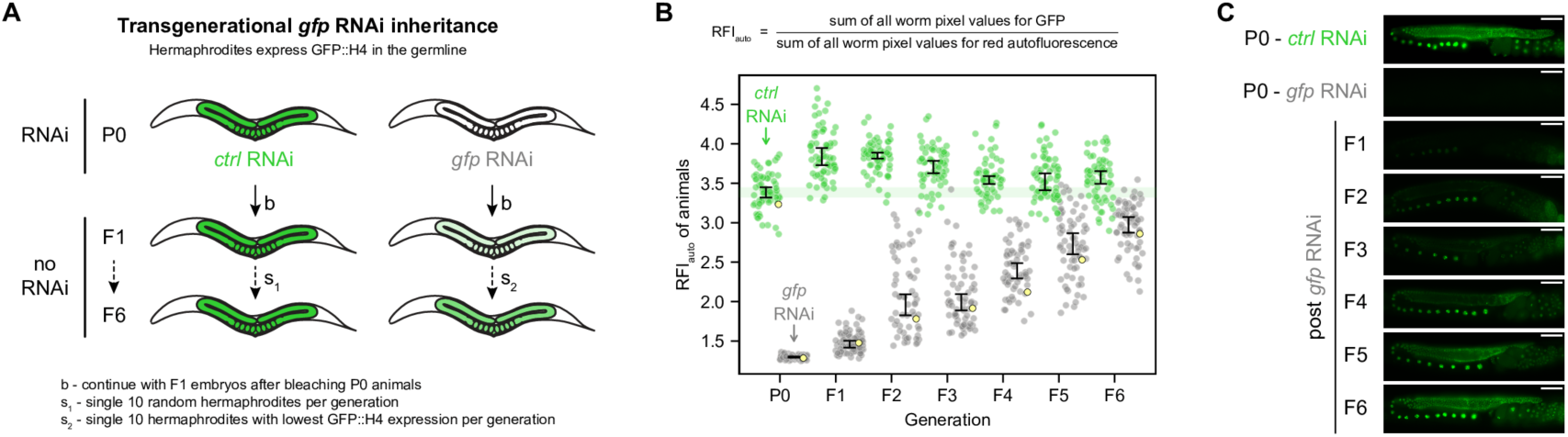
Transgenerational *gfp* RNAi inheritance of GFP::H4. A, Schematic representation summarizing the effects of transgenerational *gfp* RNAi inheritance in hermaphrodites expressing GFP::H4 in the germline. Germline color illustrates GFP::H4 expression: green – expressed, shades of light green – expressed at lower levels, white – silenced. B, Relative fluorescence intensity (RFIauto) of GFP::H4 in hermaphrodites before and during transgenerational RNAi inheritance. Only P0 hermaphrodites were treated with either *control* RNAi (green) or *gfp* RNAi (grey). Each dot represents an individual animal, with yellow dots referring to representative micrographs shown in (C). 95 % confidence intervals of the median are shown as black error bars for all samples as well as green line for the P0 generation treated with *control* RNAi. Sample sizes: P0 = ∼ 60 animals per condition, F1 – F6 = ∼ 60 animals from 10 founders per condition. C, Widefield fluorescence micrographs of representative hermaphrodites before and during transgenerational *gfp* RNAi inheritance, as indicated in (B). GFP::H4 appears in green. Generations and RNAi conditions are indicated to the left of the micrographs. Scale bars: 50 µm.

**Appendix Figure S2.**
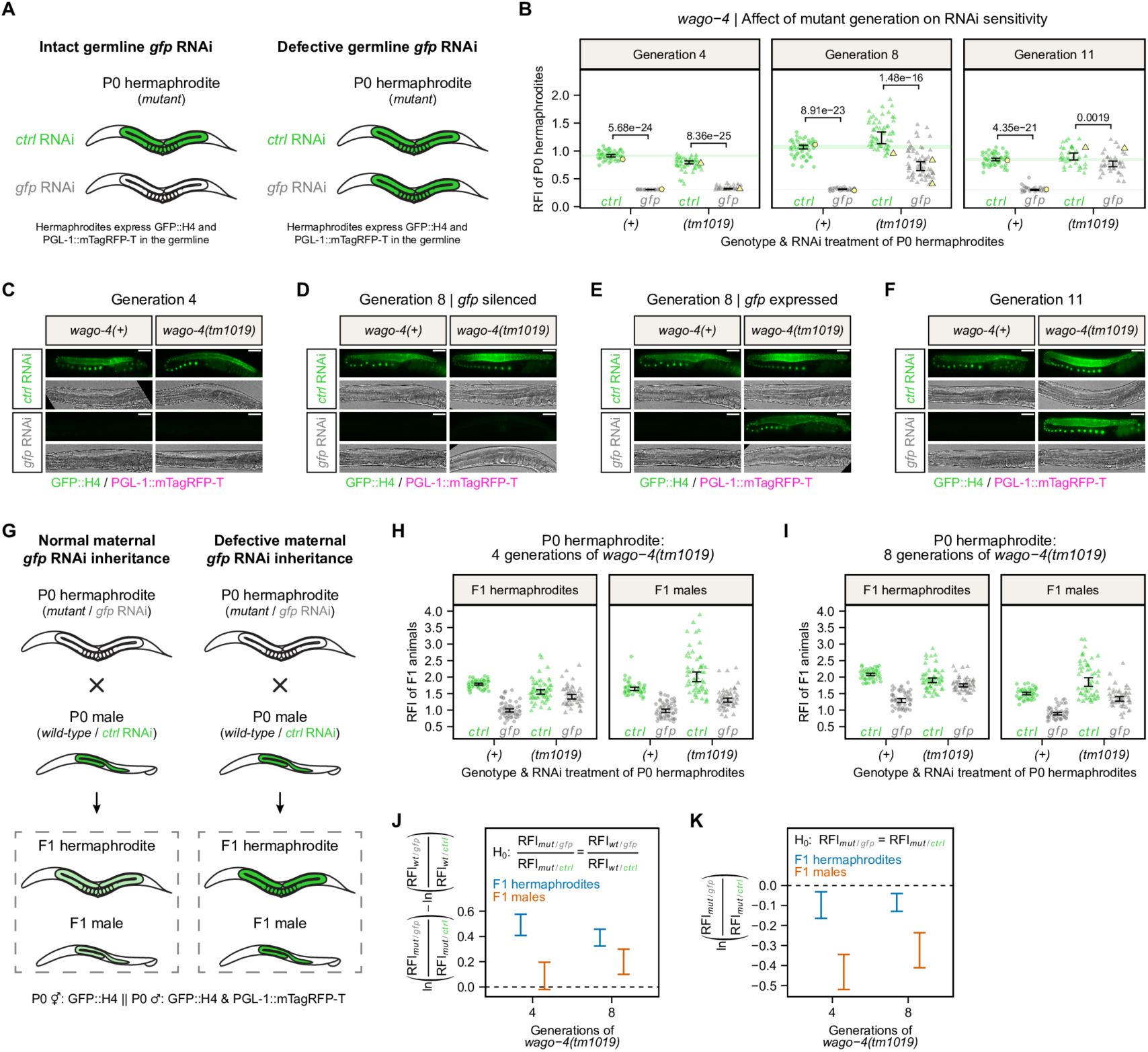
*wago-4* mutant hermaphrodites become transgenerationally insensitive to germline RNAi. A, Schematic representations summarizing the *gfp* RNAi effect on mutant P0 hermaphrodites expressing GFP::H4 and PGL-1::mTagRFP-T in the germline. Germline color illustrates GFP::H4 expression: green – expressed, white – silenced. B, Relative fluorescence intensity (RFI) of GFP::H4 in P0 hermaphrodites treated with either *control* RNAi (green) or *gfp* RNAi (grey). P0 hermaphrodites were either wild-type (circle) or mutant (triangle) for *wago-4*, with latter being analyzed after 4, 8 and 11 generations of mutant homozygosity. Each dot represents an individual animal, with yellow dots referring to representative micrographs shown in (C-F). 95 % confidence intervals of the median are shown as black error bars for all samples as well as green (*control* RNAi) and grey (*gfp* RNAi) lines for wild-type conditions. Raw data of ‘Generation 8’ is identical to Figure 2E. Sample size = ∼ 60 animals per condition. (exception: ∼ 40 *wago-4(tm1019)* animals for ‘Generation 11’). C-F, Widefield fluorescence micrographs of representative P0 hermaphrodites treated with either *control* RNAi or *gfp* RNAi, as indicated in (B). P0 hermaphrodites were either wild-type or mutant for *wago-4*, with latter being analyzed after 4 (C), 8 (D-E) and 11 (F) generations of mutant homozygosity. GFP::H4 and PGL-1::mTagRFP-T appear in green and magenta, respectively. Scale bars: 50 µm. G, Crossing schemes summarizing how mutations in P0 hermaphrodites affect germline GFP::H4 expression in F1 animals after maternal *gfp* RNAi inheritance. Germline color illustrates GFP::H4 expression: green – expressed, light green – expressed at lower level, white – silenced. H-I, Relative fluorescence intensity (RFI) of GFP::H4 in F1 hermaphrodites and F1 males after maternal RNAi inheritance. P0 hermaphrodites were treated with either *control* RNAi (green) or *gfp* RNAi (grey), and were either wild-type (circle) or mutant (triangle) for *wago-4*. Mutant P0 hermaphrodites carried the *wago-4(tm1019)* mutation for either 4 (H) or 8 (I) generations. P0 males were always wild-type for *wago-4* and treated with *control* RNAi. Each dot represents an individual animal. 95 % confidence intervals of the mean are shown as black error bars. Raw data of (I) is identical to Figure 3H. Sample size = ∼ 60 F1 animals from 5 P0 founders per condition. J, Comparison of the relative GFP::H4 fluorescence reduction after maternal *gfp* RNAi inheritance between F1 animals sired by *wago-4(+)* and *wago-4(tm1019)* P0 hermaphrodites. The plot summarizes the Gaussian models fitted in (H-I) and depicts 95 % confidence intervals (CIs) of differences of log fold changes. The null hypothesis (H0) expresses equality of relative GFP::H4 fluorescence reduction between wild-type and mutant condition, meaning that the mutation does not cause an enhanced or defective maternal *gfp* RNAi inheritance. A 95 % CI not including zero, is equivalent to a rejection of the null hypothesis at the 5% significance level and indicates that the mutation caused a defective (95 % CI > 0) maternal *gfp* RNAi inheritance. Color of CIs indicates sex of F1 animals: blue – hermaphrodite, red – male. K, Comparison of the relative GFP::H4 fluorescence intensity after maternal RNAi inheritance between F1 animals sired by *wago-4(tm1019)* P0 hermaphrodites treated with either *control* RNAi or *gfp* RNAi. The plot summarizes the Gaussian models fitted in (H-I) and depicts 95 % confidence intervals (CIs) of RFI log fold changes. The null hypothesis (H0) expresses equality of relative GFP::H4 fluorescence intensity between *control* RNAi and *gfp* RNAi treatments, meaning that the mutation causes a completely defective maternal *gfp* RNAi inheritance. A 95 % CI not including zero is equivalent to a rejection of the null hypothesis at the 5% significance level and indicates that the mutation does not cause a completely defective maternal *gfp* RNAi inheritance. Color of CIs indicates sex of F1 animals: blue – hermaphrodite, red – male.

**Appendix Figure S3.**
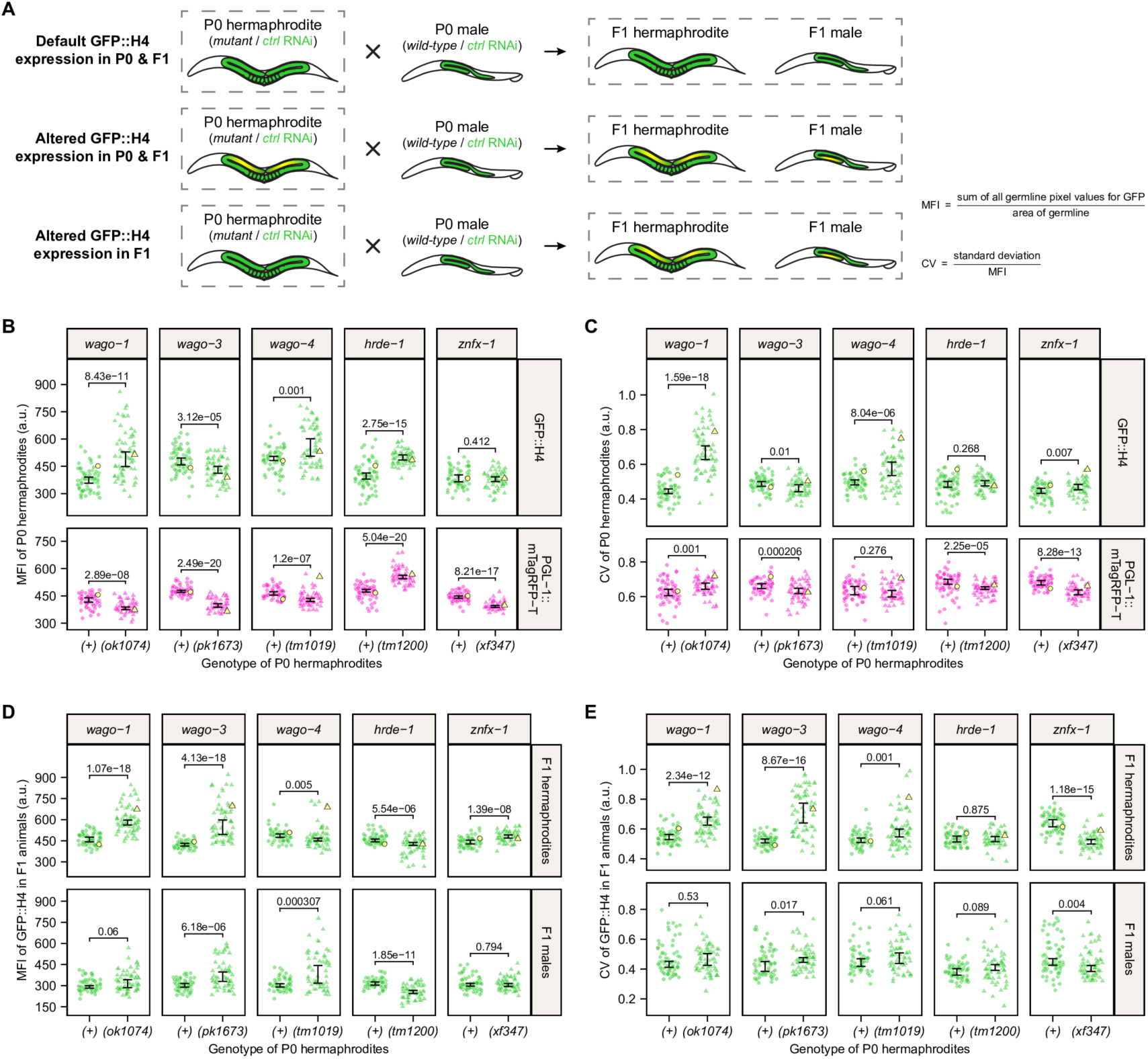
Global and local effects of mutant RNAi factors on germline GFP::H4 expression in hermaphrodites and their progeny. A, Crossing schemes summarizing how mutations in P0 hermaphrodites affect germline GFP::H4 expression in F1 animals produced by allogamy. Germline color illustrates GFP::H4 expression: uniformly green – default expression, yellow-green – mutant expression. B-C, Mean fluorescence intensity (MFI) (B) and coefficient of variation (CV) (C) of GFP::H4 (green) and PGL-1::mTagRFP-T (magenta) in P0 hermaphrodites treated with *control* RNAi. P0 hermaphrodites were either wild-type (circle) or mutant (triangle) for indicated genes. Each dot represents an individual animal, with yellow dots referring to representative animals shown in Figure 2. 95 % confidence intervals of the median are shown as black error bars. Statistically significant differences were determined using two-sided Wilcoxon rank-sum tests. Raw data is identical to Figure 2. The CV is defined as the ratio of the standard deviation to the mean. Sample size = ∼ 60 animals per condition. D-E, Mean fluorescence intensity (MFI) (D) and coefficient of variation (CV) (E) of GFP::H4 in F1 hermaphrodites and F1 males after maternal RNAi inheritance. P0 hermaphrodites were treated with *control* RNAi, and were either wild-type (circle) or mutant (triangle) for indicated genes. P0 males were always wild-type for these genes and treated with *control* RNAi. Each dot represents an individual animal, with yellow dots referring to representative animals shown in Figure 3. 95 % confidence intervals of the median are shown as black error bars. Statistically significant differences were determined using two-sided Wilcoxon rank-sum tests. Raw data is identical to Figure 3. The CV is defined as the ratio of the standard deviation to the mean. Sample size = ∼ 60 F1 animals from 5 P0 founders per condition.

**Appendix Figure S4.**
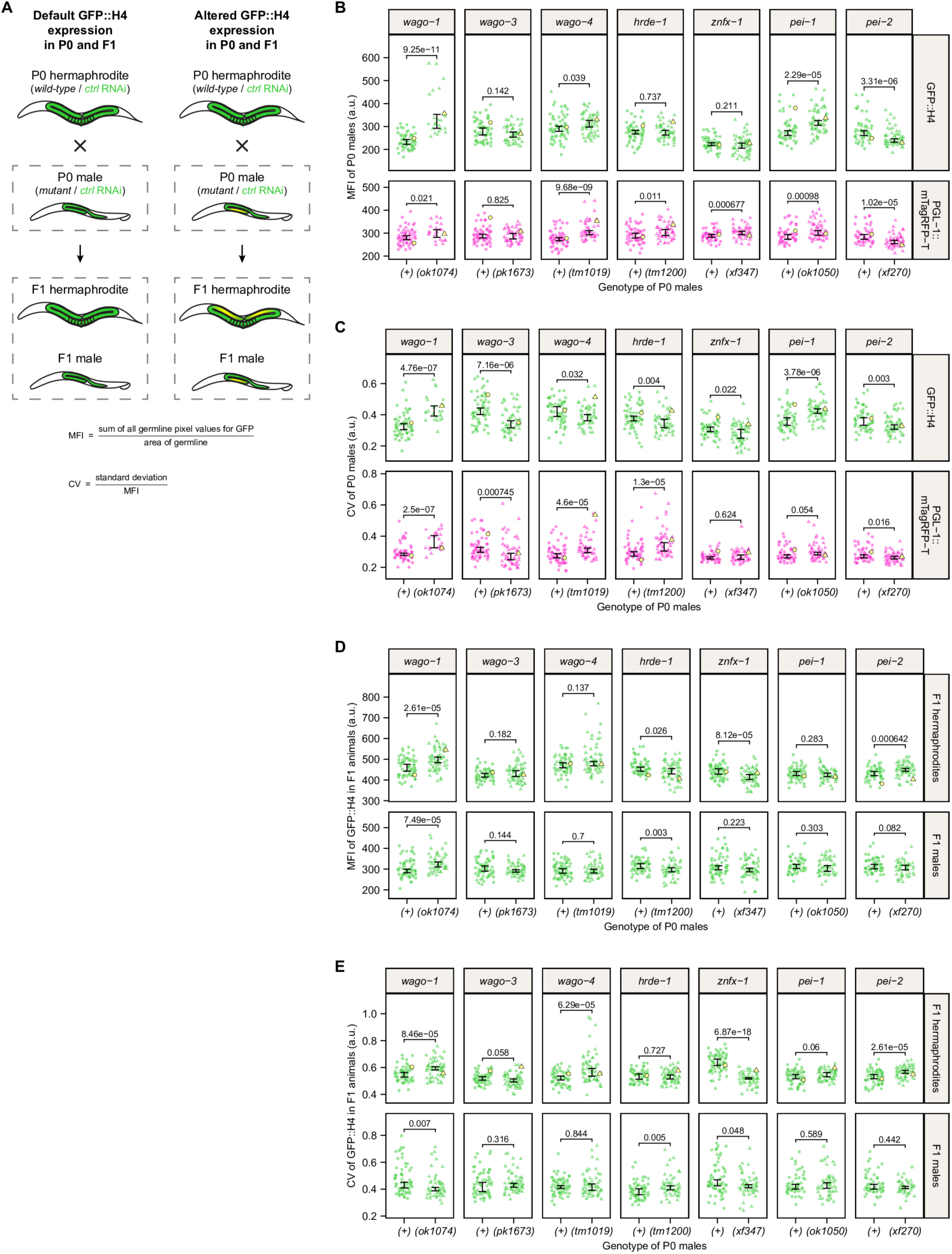
Global and local effects of mutant RNAi factors on germline GFP::H4 expression in males and their progeny. A, Crossing schemes summarizing how mutations in P0 hermaphrodites affect germline GFP::H4 expression in F1 animals produced by allogamy. Germline color illustrates GFP::H4 expression: uniformly green – default expression, yellow-green – mutant expression. B-C, Mean fluorescence intensity (MFI) (B) and coefficient of variation (CV) (C) of GFP::H4 (green) and PGL-1::mTagRFP-T (magenta) in P0 males treated with *control* RNAi. P0 males were either wild-type (circle) or mutant (triangle) for indicated genes. Each dot represents an individual animal, with yellow dots referring to representative animals shown in Figure 4. 95 % confidence intervals of the median are shown as black error bars. Statistically significant differences were determined using two-sided Wilcoxon rank-sum tests. Raw data is identical to Figure 4. The CV is defined as the ratio of the standard deviation to the mean. Sample size = ∼ 60 animals per condition (exception: ∼ 23 *wago-1(ok1074)* animals). D-E, Mean fluorescence intensity (MFI) (D) and coefficient of variation (CV) (E) of GFP::H4 in F1 hermaphrodites and F1 males after paternal RNAi inheritance. P0 males were treated with *control* RNAi, and were either wild-type (circle) or mutant (triangle) for indicated genes. P0 hermaphrodites were always wild-type for these genes and treated with *control* RNAi. Each dot represents an individual animal, with yellow dots referring to representative animals shown in Figure 5. 95 % confidence intervals of the median are shown as black error bars. Statistically significant differences were determined using two-sided Wilcoxon rank-sum tests. Raw data is identical to Figure 5. The CV is defined as the ratio of the standard deviation to the mean. Sample size = ∼ 60 F1 animals from 5 P0 founders per condition.

## TABLES AND THEIR LEGENDS

**Supplementary Table 1.** Strains used in this study.

| Strain | Genotype | Source |
| --- | --- | --- |
| N2 | *C. elegans* wild isolate var Bristol | CGC<sup>1</sup> |
| RB1096 | *wago-1(ok1074)* I. | CGC<sup>1</sup> |
| NL5117 | *wago-3(pk1673)* I. | CGC<sup>1</sup> |
| RFK22 | *wago-4(tm1019)* II. | (Barstead *et al*, 2012) |
| YY538 | *hrde-1(tm1200)* III. | CGC<sup>1</sup> |
| RB1083 | *pei-1(ok1050)* IV. | CGC<sup>1</sup> |
| RFK1342 | *pei-2(xf270)* V. | (Schreier *et al*, 2022) |
| RFK1086 | *pgl-1(xf233[pgl-1::mTagRfp-t])* IV. | (Schreier *et al*, 2022) |
| EG6699 | *ttTi5605* II; *unc-119(ed3)* III; *oxEx1578[eft-3p::gfp + Cbr-unc-119(+)]*. | (Frøkjær-Jensen *et al*, 2012b) |
| RFK1305 | *xfSi255[his-67p::gfp::his-67::tbb-2 3'UTR + Cbr-unc-119(+)]* II. | This study |
| RFK1622 | *xfSi255[his-67p::gfp::his-67::tbb-2 3'UTR + Cbr-unc-119(+)]* II; *him-5(e1490)* V. | This study |
| RFK1527 | *xfSi255[his-67p::gfp::his-67::tbb-2 3'UTR + Cbr-unc-119(+)]* II; *pgl-1(xf233[pgl-1::mTagRfp-t])* IV. | This study |
| RFK1513 | *xfSi255[his-67p::gfp::his-67::tbb-2 3'UTR + Cbr-unc-119(+)]* II; *wago-1(ok1074)* I. | This study |
| RFK1520 | *xfSi255[his-67p::gfp::his-67::tbb-2 3'UTR + Cbr-unc-119(+)]* II; *wago-3(pk1673)* I. | This study |
| RFK1515 | *xfSi255[his-67p::gfp::his-67::tbb-2 3'UTR + Cbr-unc-119(+)]* II; *wago-4(tm1019)* II. | This study |
| RFK1494 | *xfSi255[his-67p::gfp::his-67::tbb-2 3'UTR + Cbr-unc-119(+)]* II; *hrde-1(tm1200)* III. | This study |
| RFK1615 | *xfSi255[his-67p::gfp::his-67::tbb-2 3'UTR + Cbr-unc-119(+)]* II; *znfx-1(xf347)* II. | This study |
| RFK1405 | *xfSi255[his-67p::gfp::his-67::tbb-2 3'UTR + Cbr-unc-119(+)]* II; *pgl-1(xf233[pgl-1::mTagRfp-t])* IV; *him-5(e1490)* V. | This study |
| RFK1512 | *xfSi255[his-67p::gfp::his-67::tbb-2 3'UTR + Cbr-unc-119(+)]* II; *pgl-1(xf233[pgl-1::mTagRfp-t])* IV; *him-5(e1490)* V; *wago-1(ok1074)* I. | This study |
| RFK1470 | *xfSi255[his-67p::gfp::his-67::tbb-2 3'UTR + Cbr-unc-119(+)]* II; *pgl-1(xf233[pgl-1::mTagRfp-t])* IV; *him-5(e1490)* V; *wago-3(pk1673)* I. | This study |
| RFK1514 | *xfSi255[his-67p::gfp::his-67::tbb-2 3'UTR + Cbr-unc-119(+)]* II; *pgl-1(xf233[pgl-1::mTagRfp-t])* IV; *him-5(e1490)* V; *wago-4(tm1019)* II. | This study |
| RFK1473 | *xfSi255[his-67p::gfp::his-67::tbb-2 3'UTR + Cbr-unc-119(+)]* II; *pgl-1(xf233[pgl-1::mTagRfp-t])* IV; *him-5(e1490)* V; *hrde-1(tm1200)* III. | This study |
| RFK1619 | *xfSi255[his-67p::gfp::his-67::tbb-2 3'UTR + Cbr-unc-119(+)] II; pgl-1(xf233[pgl-1::mTagRfp-t]) IV; him-5(e1490) V; znfx-1(xf347) II.* | This study |
| RFK1471 | *xfSi255[his-67p::gfp::his-67::tbb-2 3'UTR + Cbr-unc-119(+)] II; pgl-1(xf233[pgl-1::mTagRfp-t]) IV; him-5(e1490) V; pei-1(ok1050) IV.* | This study |
| RFK1472 | *xfSi255[his-67p::gfp::his-67::tbb-2 3'UTR + Cbr-unc-119(+)] II; pgl-1(xf233[pgl-1::mTagRfp-t]) IV; him-5(e1490) V; pei-2(xf270) V.* | This study |

<sup>1</sup> *Caenorhabditis* Genetics Center

**Supplementary Table 2.** Plasmids used in this study.

| Plasmid | Description | Source |
| --- | --- | --- |
| pCFJ601 | Induces Mos1 transposition by injection. | Addgene |
| pMA122 | Negative selection, heat-shock inducible. | Addgene |
| pGH8 | Visual extra-chromosomal array marker. Red, nervous system, cytosolic. | Addgene |
| pCFJ104 | Visual extra-chromosomal array marker. Red, body wall muscle, cytosolic. | Addgene |
| pCFJ90 | Visual extra-chromosomal array marker. Red, pharynx, cytosolic. | Addgene |
| pCFJ350 | Multiple cloning site vector for MosSCI on locus *ttTi5605* | Addgene |
| pDD282 | *gfp::SEC::3xflag* parental vector for CRISPR/Cas9 insertions | Addgene |
| pRFK4197 | [*his-67p::gfp::his-67::tbb-2* 3'UTR] insertion in pCFJ350 | This study |
| pRFK2411 | Expression of Cas9 and sgRNA(F+E) without protospacer | (Schreier *et al*, 2022) |
| pRFK2412 | Expression of sgRNA(F+E) without protospacer | (Schreier *et al*, 2022) |
| pRFK2588 | Expression of Cas9 and *unc-58* sgRNA1(F+E) | (Schreier *et al*, 2022) |
| pRFK3358 | Expression of *znfx-1* sgRNA1(F+E) | This study |
| pRFK3359 | Expression of *znfx-1* sgRNA2(F+E) | This study |
| pRFK3360 | Expression of *znfx-1* sgRNA3(F+E) | This study |
| pRFK3361 | Expression of *znfx-1* sgRNA4(F+E) | This study |
| L4440 | Empty backbone for RNAi feeding experiments | Addgene |
| pRK4103 | *gfp* sequence from pDD282 inserted in L4440 | This study |

**Supplementary Table 3.**
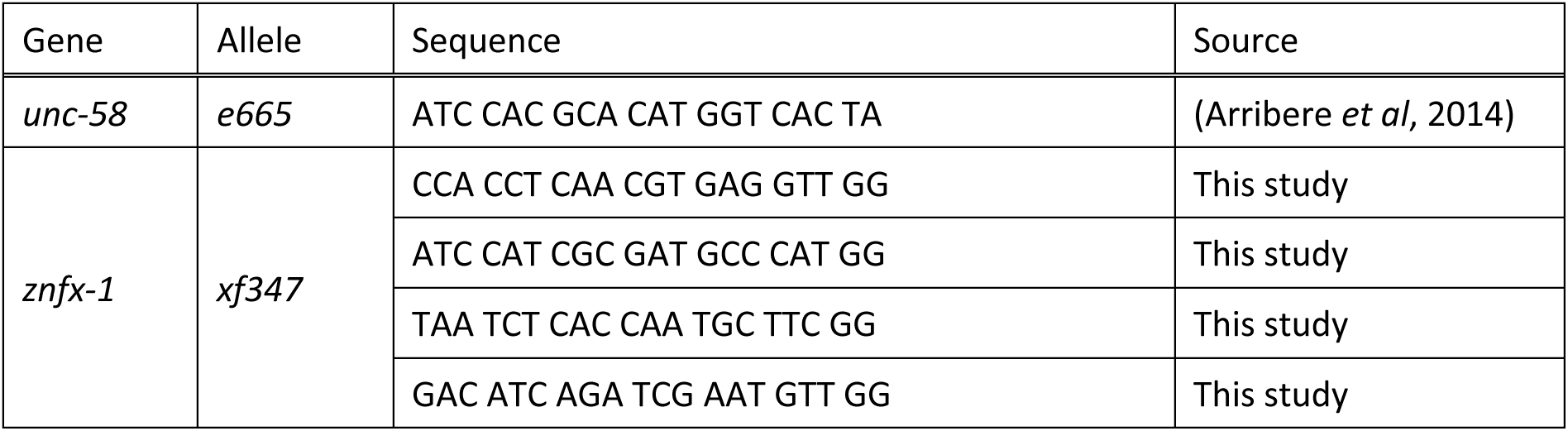
Protospacer sequences used in this study.

